# Common insecticide affects spatial navigation in bats at environmentally-realistic doses

**DOI:** 10.1101/2022.09.14.508021

**Authors:** Natalia Sandoval-Herrera, Linda Lara-Jacobo, Juan S. Vargas Soto, Paul A. Faure, Denina Simmons, Kenneth Welch

## Abstract

Bats are potentially exposed to pesticides via foraging in croplands. Common pesticides like organophosphates are neurotoxic for vertebrates and even low doses can impair essential processes such as locomotion and cognition. These sublethal effects are usually studied using molecular biomarkers with limited ecological relevance. Behavioral studies, in contrast, represent a more informative yet sensitive approach. Spatial navigation, for example, is an ecologically relevant behavior that is modulated by cellular pathways potentially targeted by neurotoxicants. We evaluated whether bats’ ability to memorize and navigate novel spaces was negatively affected by environmental relevant doses of chlorpyrifos, a common organophosphate insecticide. We also tested how the behavioral response correlated with molecular biomarkers. We orally dosed captive big brown bats (*Eptesicus fuscus*) with chlorpyrifos and studied exploratory behavior in two testing arenas. We evaluated similarity of stereotype flight trajectories in a flight tent, and associative memory in a Y-maze. We quantified brain cholinesterase (ChE) activity as a cellular biomarker and employed non-targeted proteomics as molecular biomarkers. Bats exposed to chlorpyrifos were less explorative and made more incorrect choices in the Y-maze, but the consistency of their flight trajectories was unaffected. Exposed bats had 30% lower ChE activity, showed down-regulation of proteins involved in memory (VP37D), learning and sound perception (NOX3). Other important nervous system processes such as synaptic function, plasticity, oxidative stress, and apoptosis were enriched in chlorpyrifos-exposed bats. These results support the sensitivity of behavior as a biomarker of toxicity and the importance of considering other levels of organization to help explain the mechanisms underlying altered behavior due to human activities.

## Introduction

Bats provide essential ecosystem services, such as arthropod suppression, and a sustained, thriving population is vital for ecosystem health [1]. Insectivorous bats, for example, are efficient bio-control agents eating large amounts of insects every night, including agricultural pests [2,3]. Foraging in croplands, however, makes bats vulnerable to pesticide exposure through consumption of contaminated prey or coming into direct contact with sprayed toxicants while flying and/or grooming [3–5]. In addition to a high risk of exposure, bats present physiological and ecological characteristics such as high metabolism, low fecundity and long lifespan that could make them especially vulnerable to pesticide effects compared to other non-target vertebrates [4]. Similarly, bat characteristic behaviors, such as echolocation and torpor, may be adversely affected by pesticide exposure and could jeopardize individual fitness. Understanding these threats to bat survival is crucial for conservation and will help to elucidate potential causes of observed population declines worldwide.

Commonly used pesticides like organophosphates (OPs) are known to be neurotoxic for non-target species including birds [6,7], fish [8,9], and humans[10,11], but little is known about their effects on bats [12,13]. Studies of OP toxicity in mammalian model organisms, like mice and rats, show impairment of cognitive, motor, sensory, and autonomic functions even at low concentrations (less than 1/50^th^ of lethal dose) [14–16]. These effects, however, may not simply be extrapolated to bats because of unique physiological adaptations associated with flight that set them apart from non-volant mammals [2]. Moreover, physiological and behavioral effects are often sublethal and therefore difficult to detect using traditional toxicological assessments. Hence, integrative approaches that include biomarkers at different levels of biological organization may lead to a better understanding of the deleterious effects of toxicants on wildlife. Behavioral biomarkers in particular can be a sensitive and ecologically relevant alternative to evaluate neurotoxic effects and their implications on animal fitness.

Navigation by flight is an ecologically-relevant behavior in bats essential to foraging, finding roosts, migration, and sometimes even reproduction [18]. Bats use multiple cognitive processes, including associative learning and spatial memory, to integrate previous experiences and environmental cues (e.g., position, shape, color) to find resources; however, cognitive processes can be impaired by OP pesticides due to their neurotoxic mode of action. A primary target of OPs is the enzyme cholinesterase (ChE) that hydrolyzes acetylcholine, a major neurotransmitter in the peripheral and central nervous systems [13]. Inhibition of ChE activity causes an accumulation of acetylcholine at cholinergic synapses, and causes increased neural activity [6,19]. Cognitive processes like spatial navigation, learning, olfaction, and visual perception are modulated by cholinergic synapses and could therefore be affected by neurotoxicant exposure, leading to functional disruption of cholinergic pathways.

Due to its specificity, quantification of ChE activity has become the gold standard biomarker of neurotoxicity for OPs exposure [20–22]. Although ChE inhibition is the primary mode of action of OPs, other processes such as oxidative stress, DNA damage, and cell death have been linked to neurochemical and neurobehavioral effects of OPs (reviewed in Tsai & Lein, 2021). A more comprehensive understanding of such mechanisms and adverse outcomes, however, requires an exhaustive biomarker analysis, which is limited by available knowledge on the pesticide and study species. Broad characterization of the brain proteome could help reveal, with high resolution, new and relevant molecular pathways of OP toxicity [24–26]. Proteomic analyses provide a large-scale overview of protein-level expression and regulation, which facilitates linking mechanistic impacts at the molecular level to impacts at the individual level, and consequently to populations and/or ecosystems [27].

To date, few studies have examined the neurotoxic effects of pesticides on bats and most have focused on a single biomarker, ChE activity. Here, we combine cellular and molecular biomarkers of exposure with behavioural biomarkers to evaluate the neurotoxic effects of an environmentally relevant dose of Chlorpyriphos (CPF), a widely used organophosphate pesticide, on the big brown bat (*Eptesicus fuscus*). To confirm neurotoxicity at the molecular level, we quantified ChE inhibition in the bat brain and analyzed changes in the brain proteome to elucidate other molecular and cellular pathways affected by CPF exposure. Based on the known mode of action, we hypothesized that CPF would cause a reduction in brain ChE activity and in turn affect multiple cholinergic-modulated processes, including cognition. Specifically, we predicted that effects at the cellular (ChE) and molecular level (protein profile) would be reflected by neurobehavioral impairment in spatial navigation tasks such as stereotyped navigation patterns and associative memory. Both behaviors are ecologically relevant for bats and their disruption could be detrimental for fitness and survival.

## Materials and methods

### Experimental animals and housing

We used big brown bats (*Eptesicus fuscus*) from a captive research colony established at McMaster University. Bats were either wild-caught in Southern Ontario or direct descendants of wild-caught individuals. Bats were housed indoors (2.5 × 1.5 × 2.3 m; l x w x h) in a mixed-sex colony where the temperature and lighting varied with ambient conditions, and bats had access to a larger outdoor flying area (2.5 × 3.8 × 2.7 m) with tree branches and hanging vines as habitat enrichment (Skrinyer et al. 2017). Bats had *ad libitum* access to water and food (yellow mealworms *Tenebrio molitor*; Reptile Feeders, Norfolk, ON), as well as natural (hollowed tree with bark) and artificial (towels) roosts. All bats selected for experiments were adult females that had lived in the colony for at least 6 months (N = 18, 6 bats per treatment). During experiments, bats were kept overnight in stainless steel wire mesh cages (28 × 22 × 18 cm, ¼” mesh) in an indoor holding room. Bats were housed individually so they could be video-monitored five days before exposure and during exposure. Behavioral and neurological changes were evaluated *in situ* at different time points. At experimental endpoint, bats were euthanized, and their tissues were extracted for biochemical analysis. All procedures were approved by the Animal Research Ethics Board of McMaster University (AUP #16-06-25 and #20-05-20) and conformed to the Guidelines for the Care and Use of Experimental Animals in Research published by the Canadian Council on Animal Care.

### Pesticide exposure

Bats (n = 6) were given an oral dose of the insecticide chlorpyrifos (CPF) mixed in oil once a day at the same time (1800 hrs) for three consecutive days (72 hrs) (i.e., CPF-3d). Pesticide application on large plantations might require more than three days for complete coverage, thus we repeated the experiment on new bats (n = 6) using the same daily oral dose but increasing the exposure to seven days (i.e., CPF-7d). In both experiments, bats were dosed by feeding them mealworms (*T. molitor)*, injected with a solution of CPF. In the control group (n = 6), bats were fed mealworms injected only with the oil vehicle (i.e., a sham treatment). Each bat was given a pesticide dose exposure of 10 μg/g of bat body weight (BW) per day. This dose falls in the lower range of concentrations that an insectivorous bat could realistically intake when foraging in croplands, and corresponds to the benchmark CPF dose (BMD10: central estimate of a 10 % increase in response) reported to alter 10% of plasma cholinesterase (ChE) activity (i.e., a neurotoxicity biomarker) in big brown bats [12]. It is also an estimate of daily intake based on the concentration of Chlorpyrifos reported from arthropods collected in crops and the approximate ingestion rate normalized by body weight [28,29].

### Behavioral tests

We evaluated three behaviors that are ecologically-relevant for free-ranging bats and logistically simple to operationalize in the laboratory: (1) stereotyped flight, (2) associative memory, and (3) righting reflex response. These behaviors represent different levels of cognitive complexity in their execution (e.g., learning *versus* reflex response), and were intended to provide a holistic understanding of the effects of CPF on different aspects of bat cognitive function. Behavioral experiments were conducted at night (~2100 hrs) three hours after treatment application. The sequence of testing was: (1) stereotyped flight, (2) associative memory (Y-maze), and (3) righting reflex.

### Stereotype exploration flight

When echolocating bats are introduced into a novel space they are reported to quickly develop individual stereotyped flight patterns, such as repetitive loops along a stable trajectory [30].We evaluated bats’ ability to navigate a new space by comparing the consistency of flight paths between CPF-exposed and sham-treated bats. Bats were flown in a tent (Coleman Instant Screen House, 11' × 11', Center Height 7'6") placed in the center of same room used for the associative memory trials. We tracked flight behavior by attaching a chemiluminescent stick (2.4 mm, 0.1 g) to the bat’s back and filming flight paths with an action camera (Go-Pro HERO10) mounted inside the tent ceiling. Flight trajectories were reconstructed offline using the OpenCV library in Python. We performed a quantitative analysis of the flight trajectory data in MATLAB by using a two-dimensional (2-D) cross-correlation between different sets of occupancy histograms made with the extracted location coordinates (Fig. 1), as described by Barchi et al. [31]. We evaluated two parameters of similarity: (I) consistency of trajectory between days (cross-correlation) and (II) consistency of trajectory within the same day/trial (auto correlation). Evaluating consistency between days provides information about improvement (learning) and familiarity with the space, whereas evaluating the consistency of laps within a trial provides information about the refinement or sharpness of the flight trajectory. In both cases, the measured parameter was the maximum correlation coefficient (Cmax).

**Figure 1.**
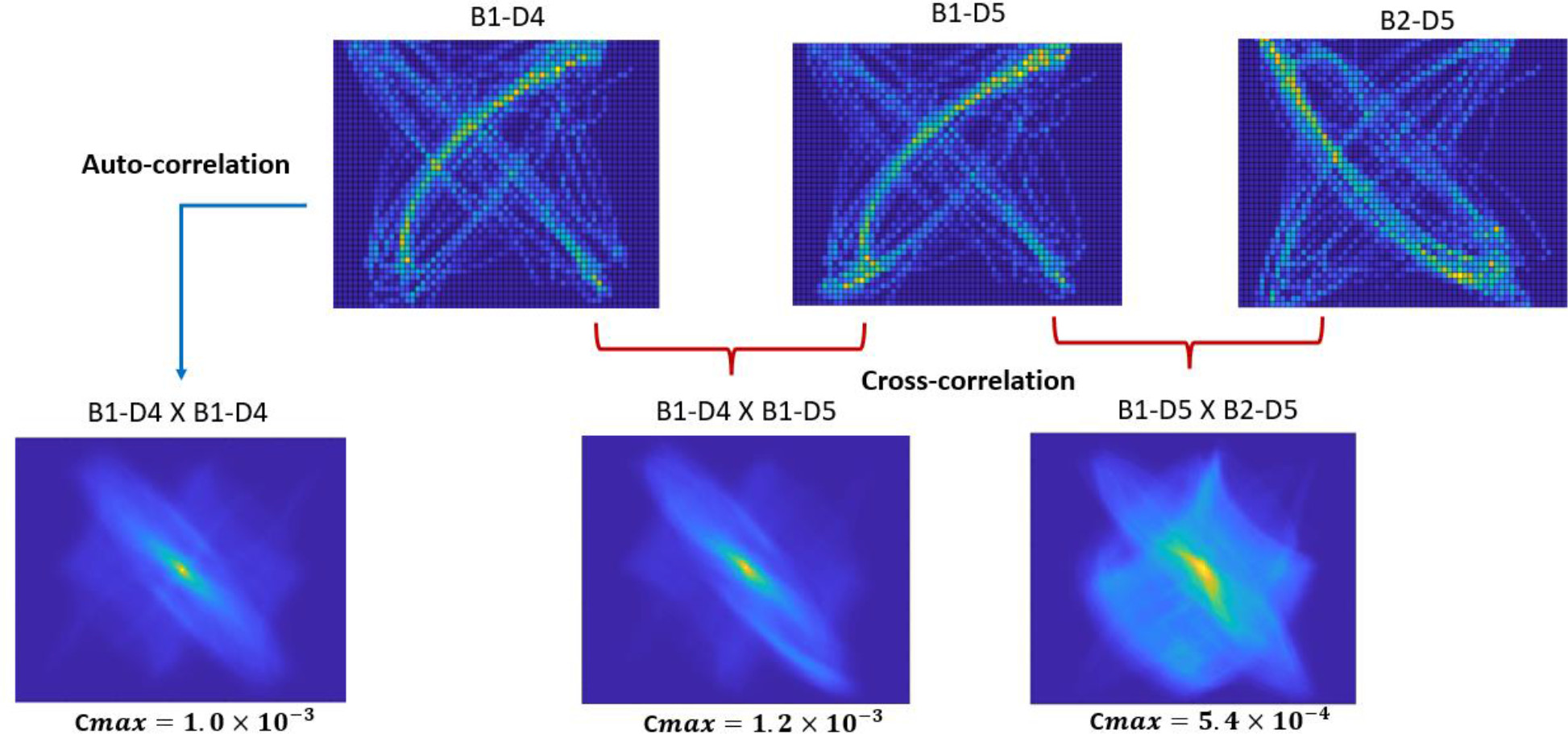
Flight path trajectory correlation analyses reveals the consistency of flight within and across trials. The three plots on the top show 2-D occupancy histograms of the reconstructed trajectory for bat 1 on day 4 (B1-D4), bat 1 on day 5 (B1-D5) and bat 2 on day 5 (B2-D5). Warmer colors (orange-yellow) indicate higher frequency of use. The three heat plots on the bottom show the auto-correlation of the path for bat 1 on day 4 (B1-D4 X B1-D4, left), the cross-correlation of paths for bat 1, days 4 and 5 (B1-D4 X B1-D5, center), and cross-correlation paths for bat 1 and bat 2 on day 5 (B1-D5 X B2-D5, right). Warmer colors indicate higher correlation values. The maximum correlation coefficient (Cmax) for each matrix comparison is printed below each plot; with higher Cmax values indicating higher similarity between the paths.

### Associative memory Y-maze

To assess associative memory, we used a 2-alternative forced choice Y-maze arena. The Y-maze is one of the simplest methods for assessing animal cognition and does not require rule learning, extensive handling or repeated manipulation [32]. The Y-maze test has been used extensively in learning and memory paradigms for rodents (Arendash et al., 2001; Conrad et al., 1997; King & Arendash, 2002; Lainiola et al., 2014; Ma et al., 2007), fish (reviewed in Cleal *et al.*, 2020) and bats [34–38]. Flight is, perhaps, the most common natural movement in bats, though many vespertilionid bats like *E*. *fuscus* are adept at crawling on surfaces when locating suitable roosts and exploring crevices and cavities [39].

Testing was conducted in the dark in a room (4.85 × 3.25 × 3.32 m) whose walls and ceiling were lined with sound-attenuating foam (Sonex® Classic; Pinta Acoustic, USA). The Y-maze walls and ceiling were constructed with cast-acrylic (Fig. 2A), and the maze was positioned at an inclination of ~30° to encourage bat’s crawling into and exploring it. An infrared camera (YI- home 1080p) placed 40 cm above the maze recorded bat exploratory behavior. At the beginning of each trial, a bat was placed in the start arm of the Y-maze and was permitted to freely move towards the response arms. Each bat received three training trails and three test trials. During training trials, one response arm of the Y-maze was blocked while the other remained open and had a reward shelter at the end (Fig.2B). The shelter was a mesh cup previously familiar to the bats (they are routinely weighed in this cup). Upon entering the mesh cup, the bat was permitted to rest in place for 5 min before it was returned to the start arm for another training trial. During the test trial, the response arm previously closed during training was opened and the bat was let free to explore. Between animals, the response arm that provided with the shelter reward was randomized.

**Figure 2.**
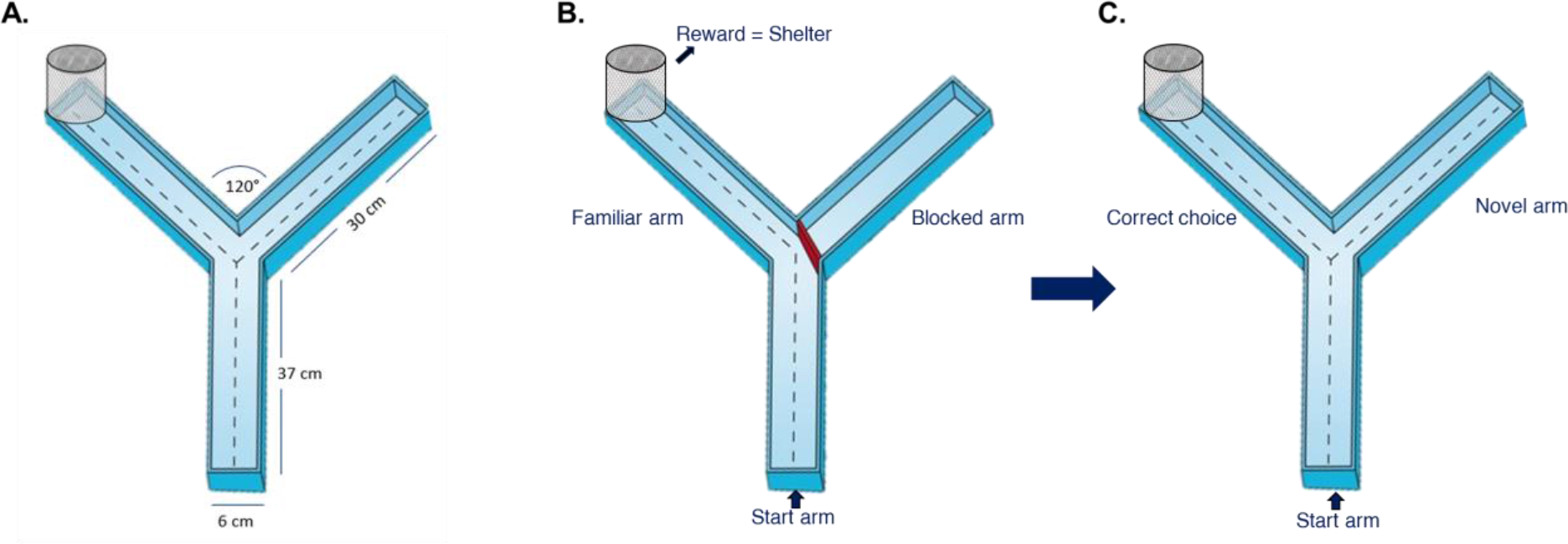
Associative memory Y-maze behavioral testing arena. A. Schematic showing dimensions of the acrylic Y-maze with a single start arm and two response arms. The dotted lines illustrate exploratory paths available to the bat. B. Y-maze set up for associative memory training trials with the right response arm blocked and the left response arm open and with a reward shelter (mesh cup) at its end; C. Y-maze set up for associative memory testing trials with both response arms open. The arm with the reward shelter is the correct choice.

During offline video analysis, the number of entries by a bat into each arm was noted as well as the crawling speed in the start arm. Bats were scored as having entered a response arm when at least two limbs were beyond the arm’s border at the Y-junction. A successful entry was defined as a bat selecting the response arm with the reward shelter (Fig.1C). Prior to analysis, the video files were renamed, and their order was randomized so the person scoring them was blind to the treatment. Due to logistical reasons the Y maze test was only conducted in bats from 2020 which were exposed to CPF for seven days (CPF-7d).

### Righting reflex response

The righting reflex test is a common behavioral assay to assess neurological function and responsiveness in rodents (Franks, 2006). It has also been used as a biomarker of loss of coordination in toxicological studies in bats (Clark 2016; Clark 2017). Rapidity of the righting reflex correlates with the state of vestibular function, coordination, and muscular strength. To examine the surface righting reflex, we placed the bat on its back on a tabletop and recorded the time taken for the animal to turn over with all four limbs on the table surface. The reflex latency was determined with offline video analysis (YI-home camera; 20 frames per second).

### Cellular and molecular biomarkers

#### Tissue collection and homogenization

Bats were euthanized by cervical dislocation ~24 hrs after receiving the final dose of CPF for their exposure regime (i.e., three or seven days). The whole brain was extracted, placed in 1.5 mL cryovials, immediately snap-frozen in dry ice and stored at −80 °C for further biochemical analysis. For testing, samples were thawed on ice, weighed, and then transferred to a new tube with ~500 μL of 0.1 M potassium phosphate buffer (pH 7.2; ≈10× the volume mass). The whole tissue and buffer were homogenized with a Branson Ultrasonics Sonifier™ SFX150 (intensity N6) while maintaining the sample on ice. One aliquot (~250 μL) was immediately used for ChE quantification, and the rest of the homogenized tissue was reserved and stored in 1.5 ml tube at −80°C for the proteomics analysis. The samples were then transported in dry ice to the Aquatic Omics Lab at the Ontario Tech University, Oshawa, Ontario.

#### Cholinesterase activity

We determined ChE activity in brain tissue using a colorimetric quantification [40], adapted to microplate by Guihermino (1996). Briefly, we used 1 mM acetylthiocholine and 0.1 mM 5,5′ dithiobis-2-dinitrobenzoic acid (DTNB) as substrate and conjugate. The reaction was measured in a full spectrum spectrophotometer (BioTek Synergy HTX) at 412 nm over 15 min. Total ChE activity was normalized to the amount of protein in the sample and expressed as U/mg of protein, U=nmol/min. Each sample was analyzed in triplicates and the average absorbance value was used. Protein quantification was conducted using the Bradford assay [41] with bovine serum albumin (BSA) as the protein standard.

### Proteomic analysis

#### Protein digestion

The homogenized sample was centrifuged for 15 mins at 14,000 x g at 4 °C and the supernatant was transferred to a new tube and purified using an Amicron Ultra10kDa centrifugal filter. We quantified protein concentration using a Bradford assay [41]. We then reduced a 50 μL volume of homogenate with the addition of 2.65 μL of 100 mM tris(2-carboxyethyl) phosphine, as a reductant, in 100 mM AB buffer, mixed using gentle vortex, and allowed to incubate at room temperature for 45 min. Proteins were then alkylated with the addition of 2.8 μL of 200 mM iodoacetamide in 100 mM AB buffer, vortexed gently, and incubated in the dark at room temperature for 45 min. At the end of the second incubation, 50 μL of chemical digestion solution (20% formic acid v/v) was added to each sample and vortexed for 5 s. Lid locks were placed on each tube and incubated at 115°C for 30 min (VWR model 96 place heating block). Samples were dried in a centrifugal evaporator (SpeedVac, Thermo-Fisher) for 40 min, stored at 4°C overnight, and then resuspended in 20 μL of a solution of 95% H_2_O, 5% acetonitrile and 0.1% formic acid. Samples were vortexed until the dried pellets were completely dissolved, and then centrifuged for 10 min at 14000 g. A 20 μL sample of the supernatant and 1 μL of internal peptide standard (H2016, Sigma-Aldrich, Oakville, ON) were added to a 2 mL screw threaded HPLC vial (Chromatographic Specialties, 12 × 32 mm) containing 250 μL PP (Polypropylene) bottom spring inserts (Canadian Life Sciences, 6 × 29 mm). Samples were stored at 4°C until instrumental analysis.

A 2 μL aliquot of the peptide solution from each sample was injected and then separated by reverse phase liquid chromatography using a Zorbax, 300SB-C18, 1.0 × 50 mm 3.5 μm column (Agilent Technologies Canada Inc., Mississauga, ON) and Agilent 1260 Infinity Binary liquid chromatographer. The Agilent 6545 Accurate-Mass Quadrupole-Time-of Flight (Q-TOF) was used as the detector in tandem to the Agilent 1200 series liquid chromatography system (see Appendix I for detailed instrumental methods).

Each analytical run included a solvent blank and a BSA digest standard (Agilent Technologies Canada Inc., Mississauga, ON) injection every 10 samples to monitor baseline, carry-over, drift, and sensitivity during the runtime. Samples were injected once per each bat.

Spectral files for each sample were analyzed using Spectrum Mill Software (Version B.04.01.141). Given the limited entries available for *E. fuscus*, peptides were searched against the Uniprot Reference Proteome of the bat family Vespertilionidae (ID 9431, 150,920 proteins; downloaded April 2021). Proteins were manually validated and accepted when at least one peptide had a score (quality of the raw match between the observed spectrum and the theoretical spectrum) greater than 5 and a %SPI (percent of the spectral intensity accounted for by theoretical fragments) greater than 60%; these are the manufacturer recommended settings for validating results with an Agilent Q-TOF mass spectrometer. After peptides were sequenced and identified by Spectrum Mill at the Tandem mass spectrometry (MS/MS) level, quantification at the precursor intensity (MS1) level was performed using the data-dependent acquisition (DDA) workflow in Skyline 20.2 (MacCoss Lab Software) with a score of 0.9, retention time window of 5 min, and 5 missed cleavages with transition settings for TOF (Pino et al., 2020).

### Statistical Analyses

For the behavioural experiments, we used a GLMM (generalized linear mixed model) approach to test the effects of treatment (CPF-exposed and sham-treated) and condition (before and after exposure) as predictors of the different cognitive tasks. To account for the repeated measures design we included the ID of the animal as a random effect. First, to evaluate the flight pattern consistency within trials (auto correlation) and between days (cross correlation) among treatments we ran GLMMs including time, treatment, and their interaction. We expected CPF exposed bats to have no improvement in consistency after being exposed compared to sham treated bats. Second, we modeled how the number of correct arm entries, total number of arm entries, crawling speed (s), and reflex latency (s) was influenced by the treatment, condition, and their interaction. Last, to examine the effect of exposure duration on the performance of the cognitive tasks, we used GLMMs including the timepoint as fixed factor across four timepoints: before exposure (BE) and one, four, and seven days after exposure (AE1, AE3, AE7), as well as treatment. In all cases, we considered an effect to be statistically significant if the coefficients 95% confidence intervals (CI) did not include zero (0). To evaluate the differences between treatments and conditions we used a Tukey pairwise post-hoc test with a Bonferroni correction. These comparisons are presented as boxplots where the line represents the median and the box represents the interquartile distance. The models were conducted using the package *lme4* [42] in R version 3.8 [43].The performance diagnostics of each model (linearity, collinearity, normality of residuals and homogeneity of variance) were evaluated using the package *performance* in R version 3.8 [43].

Proteomics data were sorted and manually consolidated on a spreadsheet (Microsoft Excel) and statistical analyses were performed with Metaboanalyst 5.0 [44]. We replaced missing values with 1/5 of the limit of detection (LOD) and normalized (using median, pareto scaling). We used the normalized data and compared the treatments with a one-way ANOVA (Fisher’s least significant difference post-hoc test with Benjamini-Hochberg false discovery rate (FDR= 0.25) and partial least squares discriminant analyses. Last, we used Revigo server to consolidate and visualize the Gene Ontology (GO) enriched pathways [45]. Revigo uses multidimensional scaling to reduce the dimensionality of a matrix of the GO terms pairwise semantic similarities. The mass spectrometry proteomics data have been deposited to the ProteomeXchange Consortium via the PRIDE partner repository with the dataset identifier PXD033247.

Finally, we used a one-way ANOVA to compare ChE activity between treatments and a Tukey HSD as post-hoc test.

## Results

### Behavioral tests

#### Stereotype exploration flight

We computed cross-correlations for the flight path trajectories of CPF-exposed and sham-treated bats for five days before and seven days after the treatment day (Fig. 3A). The maximum auto-correlation for both sham and CPF-exposed bats increased with time suggesting an increase within trial consistency in flight trajectories (β = 2.31×10-5; CI [1.11 ×10^−6^ : 4.49 ×10^−5^]; Fig. 3A), however there was no difference among treatments (β = −3.11×10^−5^; CI [−8.45 ×10-5 : 2.14 ×10^−5^]). Likewise, the maximum cross correlation between consecutive days for a given individual seemed to increase with time for both treatments (β = 3.41×10^−5^; CI [1.46 ×10^−5^ : 5.37 ×10−^5^]; Fig. 3B), however this effect was not significantly different among treatments (β = 5.85×10^−6^; CI [−4.73 ×10^−5^ : 5.89 ×10^−5^]).

**Figure 3.**
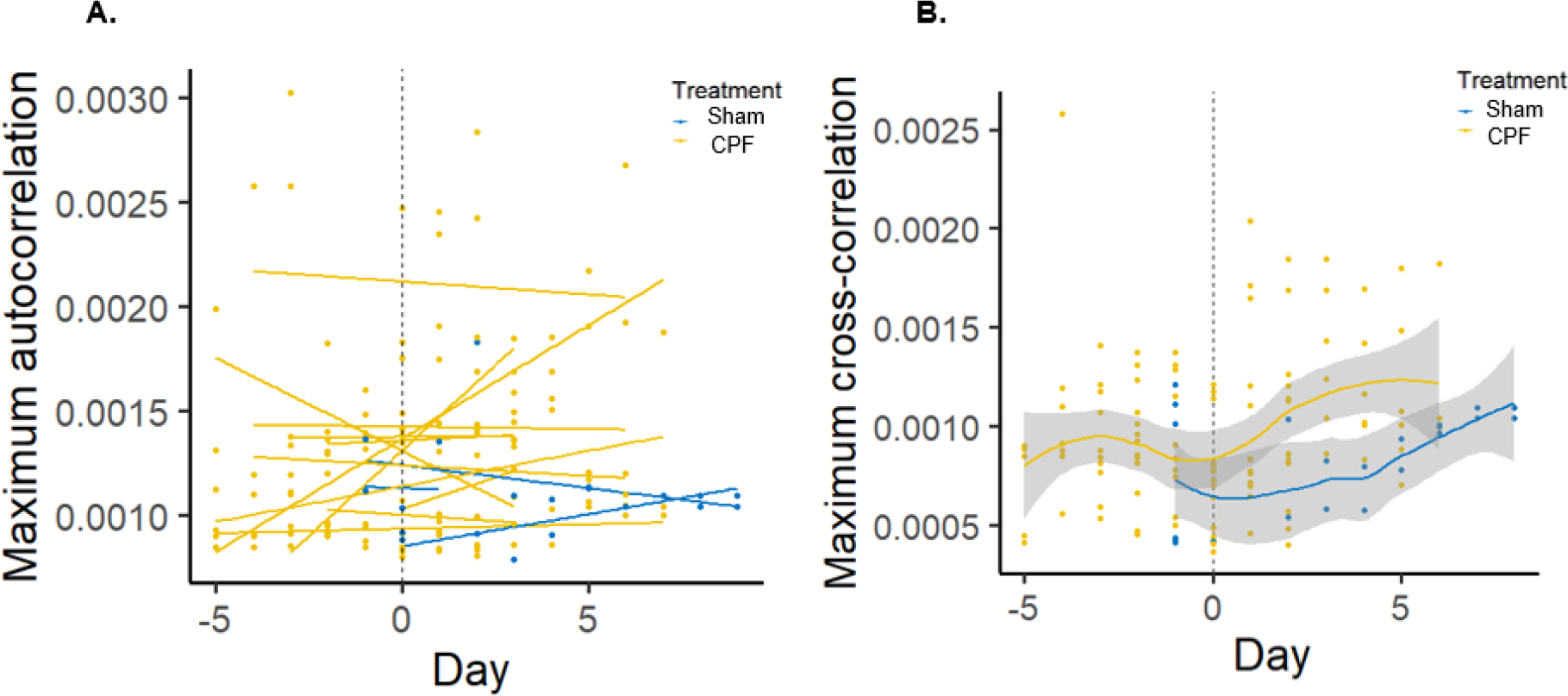
Similarity of flight trajectories in *E. fuscus* for 5 consecutive days before and after treatment in CPF-exposed and sham-treated bats. A. Comparison of maximum auto-correlation coefficient for flight laps within one trial; B. Comparison of maximum cross-correlation coefficient among consecutive days. The *dotted line* at day 0 indicates when CPF or sham-treatment dosing occurred; negative numbers represent the number of days before exposure and positive numbers are the number of days after exposure.

### Associative memory Y-maze testing

We found a significant difference in the proportion of correct entries in the Y maze arena between the two treatment exposure conditions, before and after exposure (β = 40.7; CI [4.46 : 76.9]). Bats exposed to CPF showed a reduction in the number of correct entries following pesticide exposure (t_25_ = −2.7; p = 0.04) whereas sham-treated bats showed no change in the number of correct entries (t_25_ = 0.73; p = 0.88).When comparing different timepoints after exposure to CPF (Fig. 4C), a pronounced decreased is observed three days after exposure (CPF-3d) with respect to three days after exposure (CPF-7d); however, none of the interaction terms were statistically significant. Similarly, the average number of arm entries, as indicator of exploratory activity, differed between CPF- exposed and sham-treated bats (β = −44.43; CI [−77.61 : −11.24]). Bats exposed to pesticide reduced the overall number of entries into the test arms after the treatment (t_25_ = −32.4; p = 0.01) whereas sham-treated bats explored the Y maze in a similar way before and after exposure to the sham treatment (t_25_ = 0.87; p = 0.82). This reduction in exploration was not significantly different between 1,3 and 7 days after exposure (Fig. 4D). The speed of crawling in the start arm did not differ between conditions for any of the treatments (β = 0.32; CI [−25.56 : 26.21]). CPF-exposed and sham-treated bats had similar median speeds (mean ± standard error) 70.2 ± 3.3 mm/s and 74.3 ± 7.2 mm/s, respectively.

**Figure 4.**
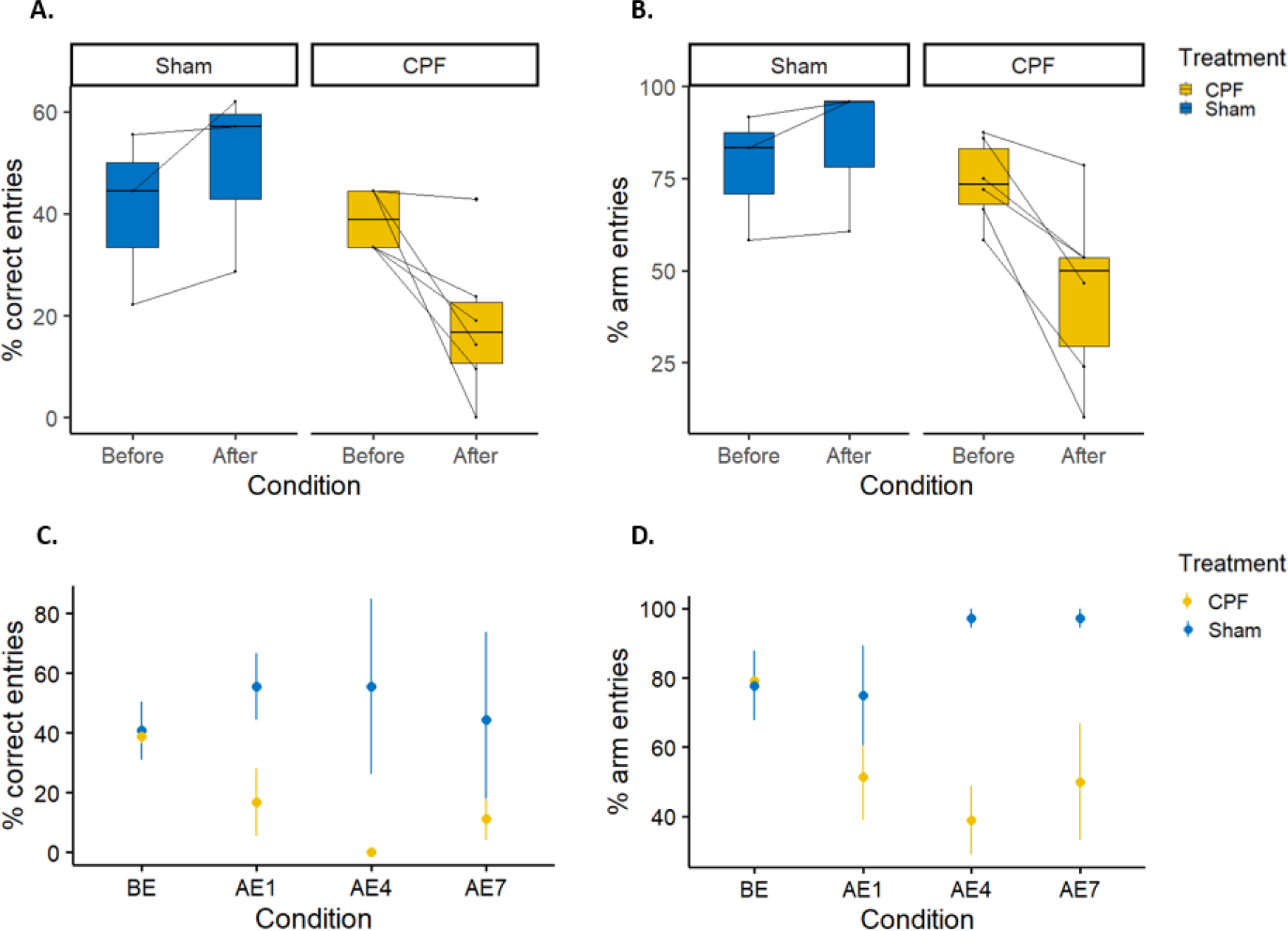
Cognitive behavior task performance of *E. fuscus* in Y-maze testing for sham-treated (blue) and bats exposed to CPF for seven days (yellow). The top row shows paired responses before and after treatment, the bottom row shows the response before exposure (BE) and 1, 3 and 7 days after exposure (i.e., AE1, AE3 and AE7); A, C. Percent correct entries to the reward arm; B, D. Percent arm entries. Thin lines in top panel connect data for the same individual.

### Righting reflex

Righting reflex latency differed between the conditions, before and after pesticide exposure, for CPF-exposed bats (β = −0.39; CI [−0.61 : −0.18]). Following oral treatment with CPF for three days, bats took 55% longer to turn over from a supine position compared to control bats in the sham-treatment (t_32.5_ = 5.26; p < 0.01; Fig. 5). Similarly, bats exposed to CPF for seven days had a significantly slower reflex response relative to sham treated bats (66% longer reflex latency; t_32.5_ = 3.94; p < 0.01).

**Figure 5.**
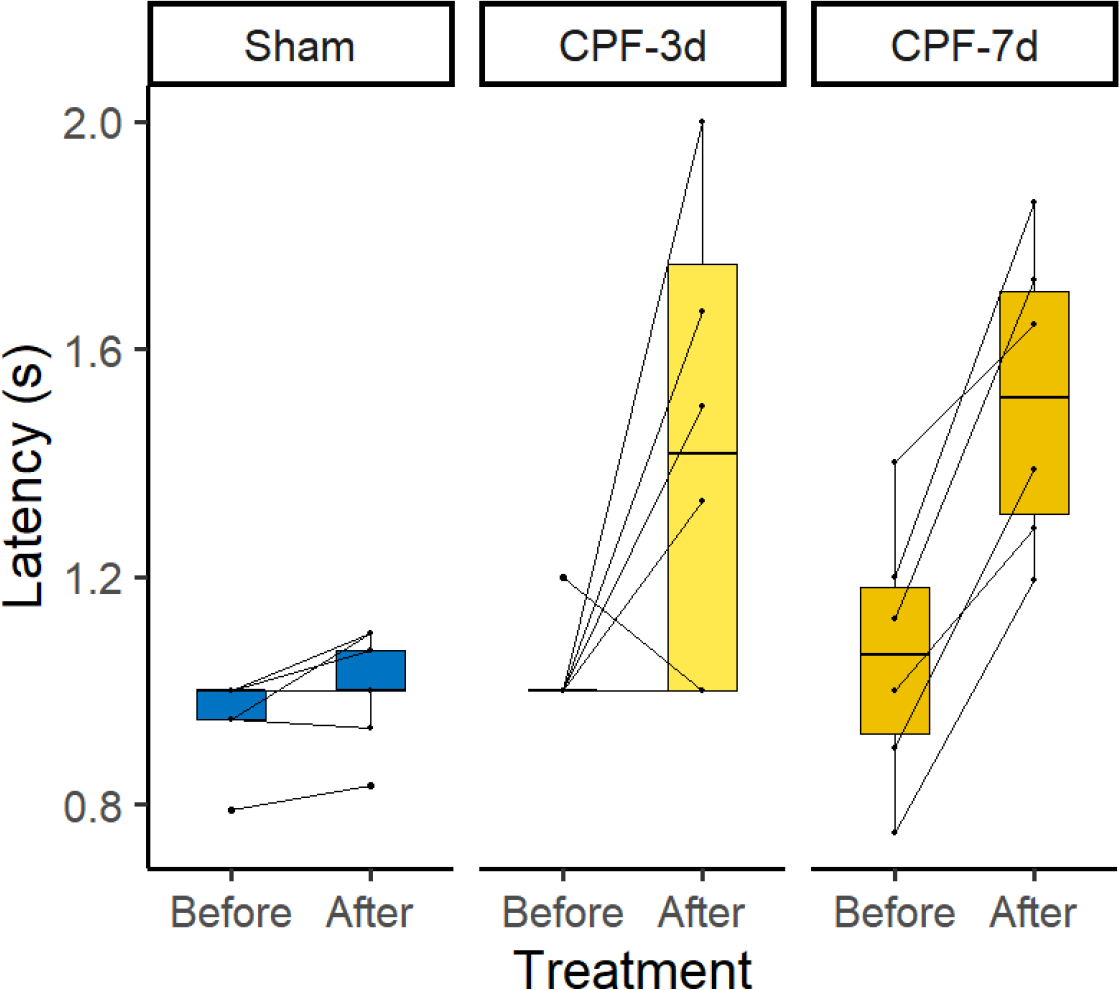
Righting reflex response in *E. fuscus* before and after exposure to a sublethal dose of chlorpyrifos (10 μL/g of BW) for three and seven consecutive days. *Thin lines* connect the reflex latency data of the same individual.

### Cholinesterase activity

Cholinesterase activity was affected by exposure to CPF (F_2,15_ = 13.84; p < 0.01; Fig. 6). Bats exposed to CPF for three days showed a 15% decrease in ChE activity compared to sham-treated bats (Tukey HSD: p < 0.05; CI = [−13.08 : −38.04]). In bats exposed for seven days, the reduction in ChE activity was 30% and significantly lower than the sham-treated bats and bats exposed to CPF for three days (Tukey HSD: p < 0.05; C.I = [−1.64 : −27.27]).

**Figure 6.**
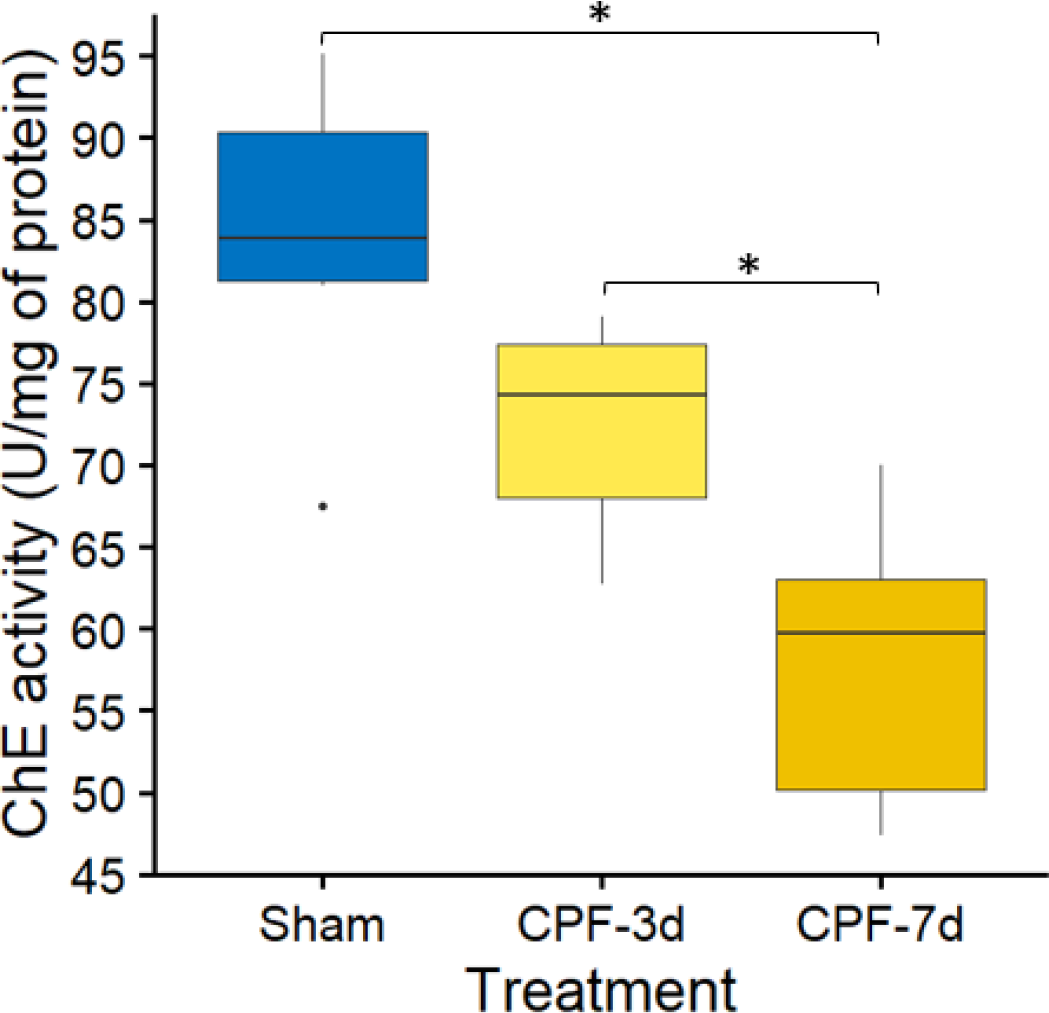
Brain cholinesterase (ChE) activity in *E. fuscus* exposed to a sublethal dose of chlorpyrifos (10 μL/g of BW) for three (CPF-3d) and seven (CPF-7d) consecutive days.

### Proteomics

We identified a total of 398 proteins in bat brain samples, most of which are highly specific to the nervous system in humans and mice (Human protein atlas, 2022). When comparing protein profiles among the treatment groups (Sham, CPF-3d and CPF-7d), we found 53 differentially abundant proteins (DAPs; P<0.1); however, the statistical significance of differential expression across the 53 proteins did not hold after a Benjamini-Hochberg correction (FDR=0.25). Our findings are nevertheless indicative of an effect with important biological implications. Overall, CPF exposure altered the abundance of proteins involved in essential biological processes such as: cell proliferation (e.g., HIVEP3, ZNF184), cellular metabolism (PTPRT, SGK1, GAA, GPATCH4), response to stimuli (SGK1, GSTA2, CYP251) and signal transduction (CRTAM, ERFE, FPR2) (Fig. 7; Supplementary Table 1).

**Figure 7.**
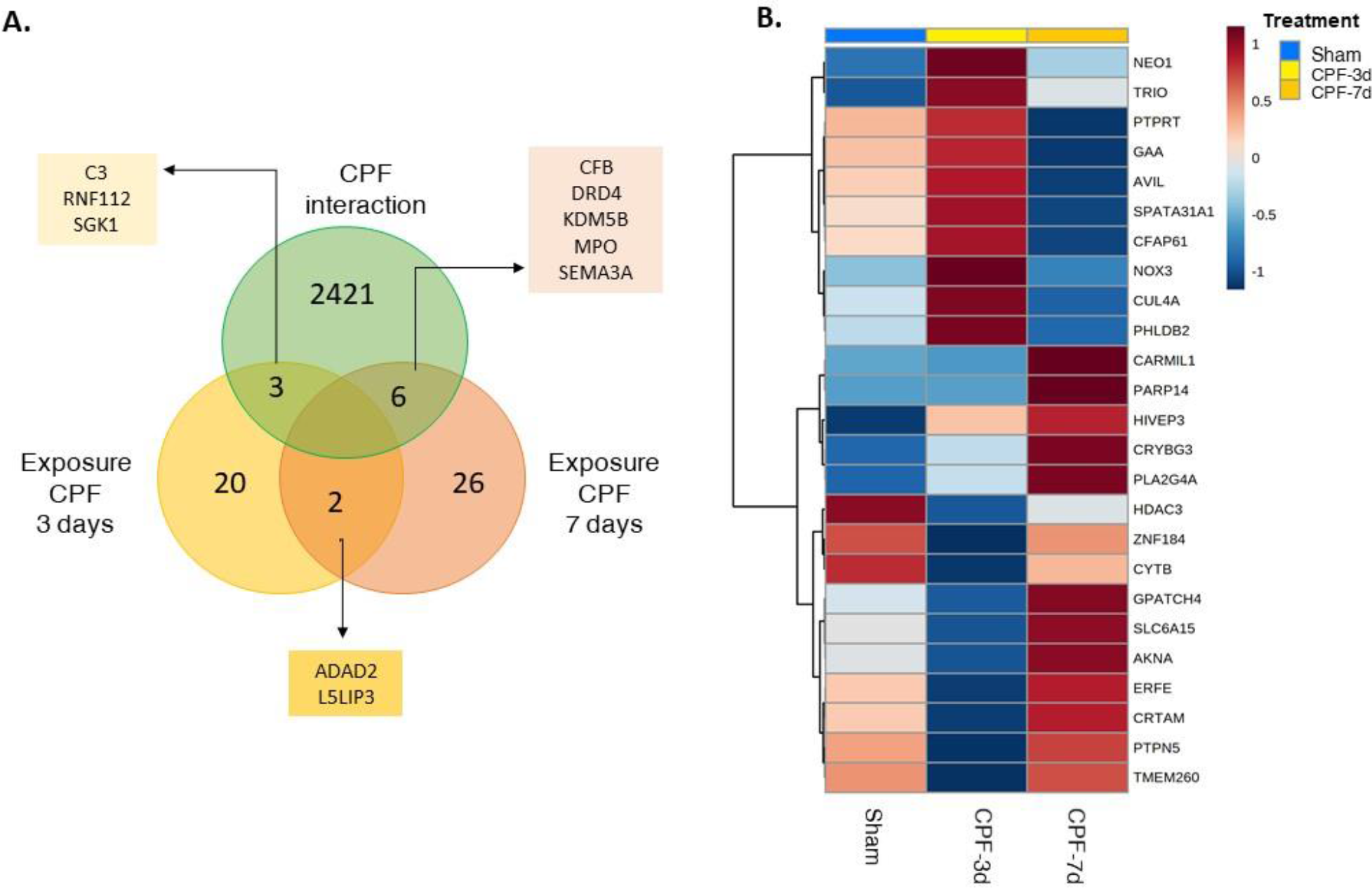
Differential expression of proteins in brain samples of *E. fuscus* exposed to the insecticide Chlorpyrifos (CPF) for three (CPF-3d) and seven (CPF-7d) consecutive days. A. Venn diagram showing the number of differentially expressed proteins identified among CPF exposure groups (*yellow* = CPF exposure for 3 days; *brown* = CPF exposure for 7 days and the number of proteins shared with the comparative toxicogenomic database for CPF (*green*). B. Clustering result shown as heatmap (Euclidean distance measure and Ward clustering algorithm) illustrating the average expression per treatment of the top 25 differentially expressed proteins (ANOVA p>0.1).

Of the DAPs, about 65% showed a uni-directional pattern of change where expression either increased or decreased as the duration of exposure increased (Fig. 8). Some proteins that increased in exposed bats participate in cellular division and apoptosis (CRYBG3, PLA2G4A), oxidative stress (PLA2G4A, CYP2S1) and inflammation (HIVEP3) processes that, if deregulated, can generate deleterious effects in nerve cells. Fewer proteins (24%) were reduced, among them, scaffold proteins like Fraser syndrome 1 (FRAS1), Serine/threonine-protein kinase (SGK1), and Receptor Protein Tyrosine Phosphatase (PTPRT). The remaining 11% of proteins showed opposite alterations in their abundance when comparing between the two exposure durations of 3 days and 7 days (Fig.7B). Among those, we detected proteins involved in DNA repair (CUL4A), cell growth rate (GPATCH4), synaptic plasticity (PTPN5) and oxidative stress (NOX3). These non-linear changes coincide with the trend observed in the associative memory evaluation where bats exposed to CPF showed the worst performance in the Y maze three days after exposure but seemed to improve after seven days of exposure (Fig.4 B).

**Figure 8.**
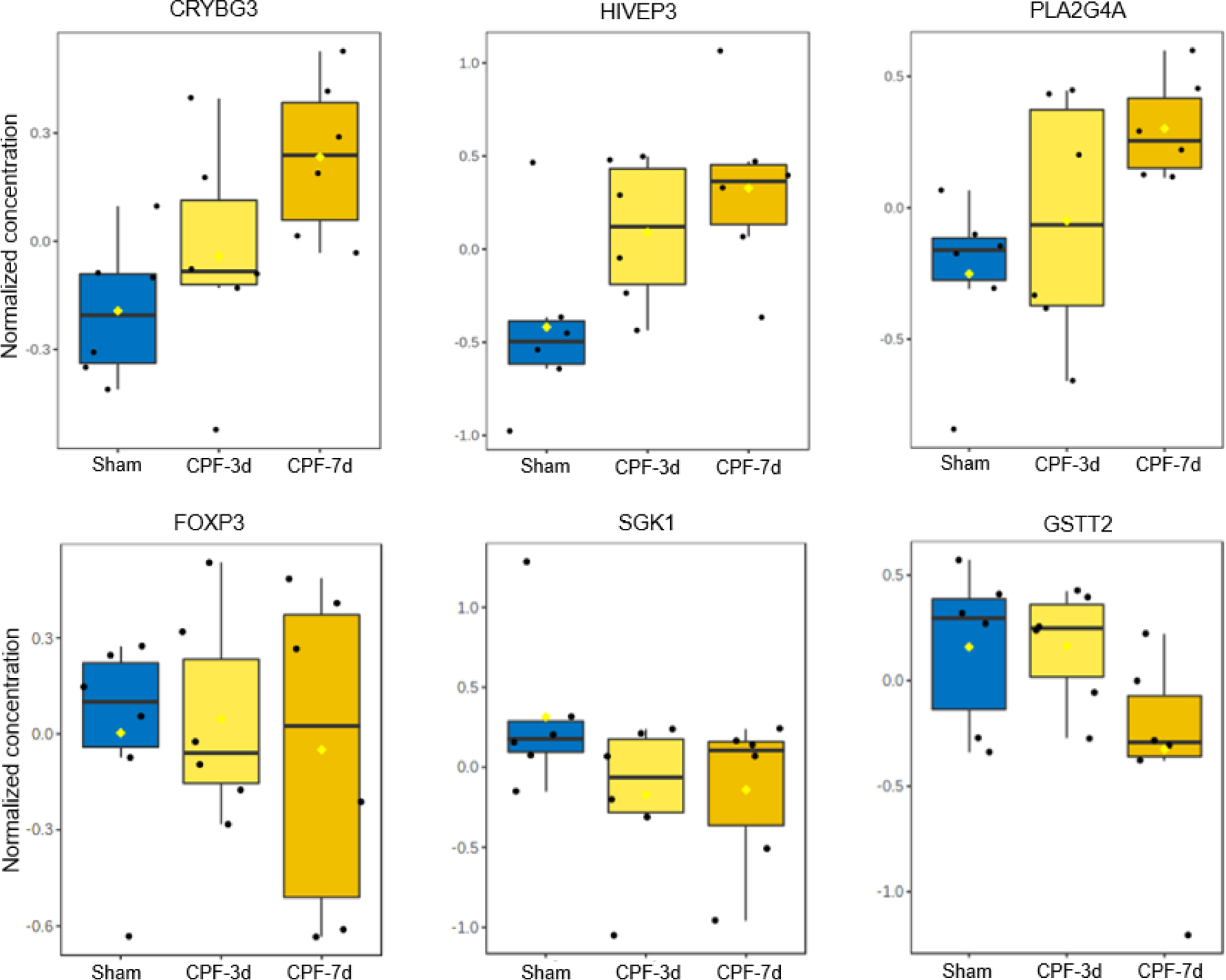
Six examples of differentially expressed proteins in brain samples of *E. fuscus* exposed to the insecticide Chlorpyrifos (CPF) for three (CPF-3d) and seven (CPF-7d) consecutive days. Proteins in the *top row* showed a positive linear response while proteins in the *bottom row* showed a negative linear response with the duration of the CPF exposure.

### Functional enrichment analysis

Gene ontology (GO) and pathway analyses revealed that the DAPs in CPF-exposed bats resulted in the enrichment of cellular processes important for nervous system function such as synaptic transmission (e.g., PTPN5, PTPRT), oxidative stress (e.g., RNF112), and apoptosis (e.g., GRIA2, PLA2G4A) (Fig.9). Proteins associated with cellular division and death were the most abundant and significantly over-represented. Pertinent to the pesticide exposure, organophosphate metabolism was one of the significantly enriched processes, with increased abundance of related proteins in CPF-exposed bats. We also found four proteins that have been previously associated with CPF toxicity according to the comparative toxicogenomic database: glutathione S-transferase theta 2 (GSTT2), syntaxin 5 (STX5), protein tyrosine phosphatase non-receptor type 5 (PTPN5 (also known as striatal-enriched protein tyrosine phosphatase [STEP]), and excitatory amino acid transporter 5 (SLC1A7) (Fig. 8.; See Table 1 for protein descriptions).

**Figure 9.**
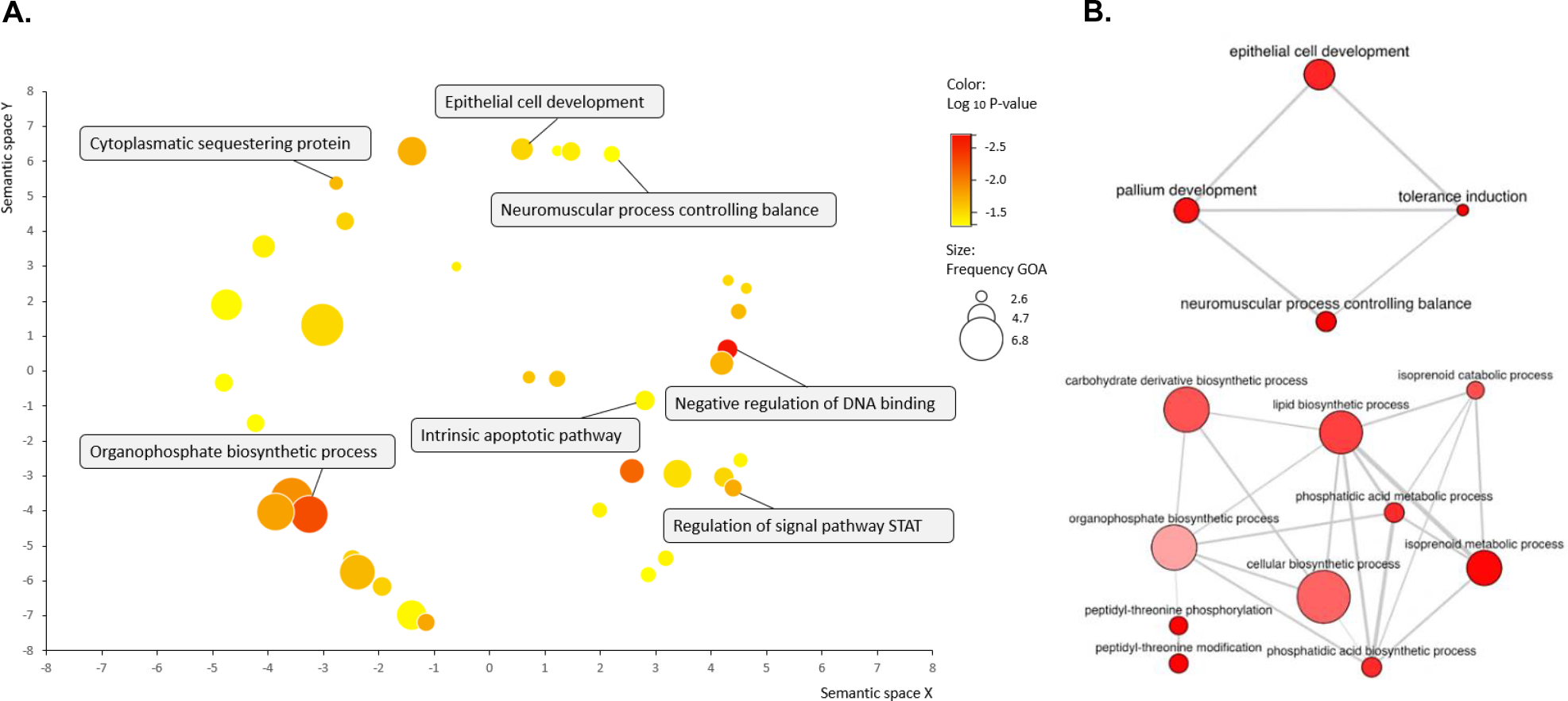
Enriched biological processes among the differentially expressed proteins in brain samples of *E. fuscus* exposed to the insecticide Chlorpyrifos (CPF). A. Consolidation of the enriched processes. The axes in panel A have no intrinsic meaning. B. Map network showing potential interactions among differentially enriched processes. In both panels, *bubble size* represents the frequency of the gene ontology (GO) term in the underlying GOA database and *bubble color* indicates level of statistical significance. Highly similar GO terms are linked by edges in the graph, where the line width indicates the degree of similarity.

**Table 1.**
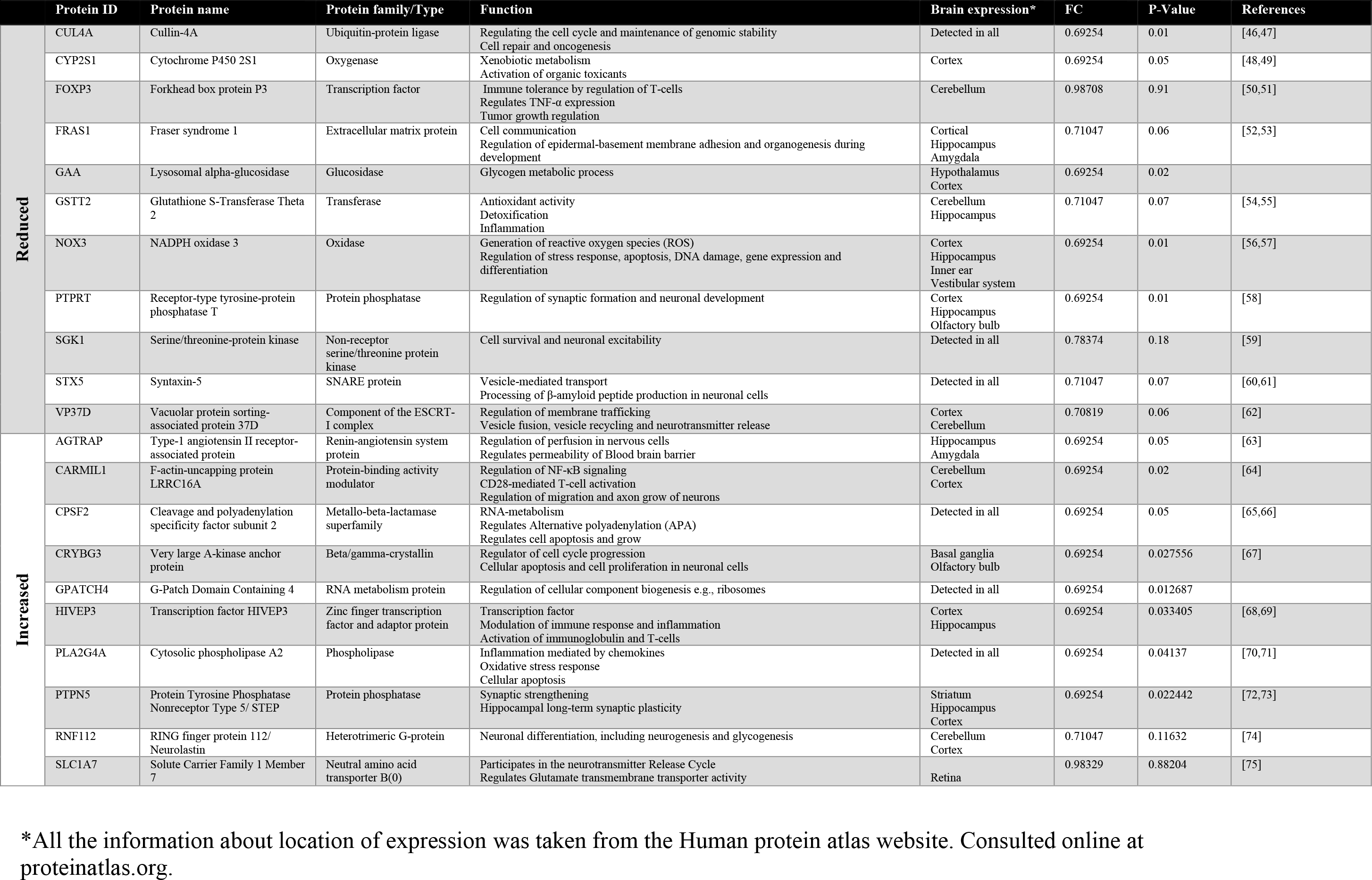
Description of relevant proteins differentially abundant in bats exposed to the insecticide Chlorpyrifos

## Discussion

Despite the critical role bats play in ecosystems coupled with decades of research into the effects of insecticides on wildlife, little is known about the risks posed to bats by insecticide use. Effects on behavior, for example, can have large implications for individual fitness and population persistence. Here we provide evidence, at the molecular and individual levels, of impairment in associative memory, which is essential for bats to effectively find food and shelter. We also found an impairment of rapidity of the righting reflex, a motor skill important for bats in maneuvers like landing. Using a proteomics approach, we detected changes in the abundance of proteins involved in neuronal function and survival, which highlights possible mechanisms underlying the observed altered behavioral phenotypes. This is the first study combining behavioral and molecular biomarkers to understand the neurotoxic effects of pesticides in a non-model mammalian species.

In several model laboratory organisms (e.g., rats, mice) there is substantial evidence that CPF neurotoxicity induces changes in behavior such as learning, fear conditioning, locomotion, and social interactions, yet few studies have reported these effects in wild/non-model vertebrates [7,12]. Most of these studies focus on ChE inhibition as the key mechanism causing disruption in behavior, and as the main proxy for OPs exposure and effects. In echolocating insectivorous bats, ChE inhibition due to OPs exposure has been reported for *Myotis lucifugus* (Clark, 1986; Clark & Rattner, 1987;) and *E. fuscus* [12], and our study found similar levels of ChE inhibition (30% for 10 μg/g) as reported previously for *E. fuscus* (40% for 10 μg/g; Eidels *et al.*, 2016) A reduction in ChE activity by more than 20% is above the reported range of natural variation in wild vertebrates, and can confidently be attributed to toxicant exposure [77]. In addition to ChE inhibition, we found enrichment of proteins involved in xenobiotic transport (TRDN), permeability of the blood brain barrier (AGTRAP), and toxicant biotransformation (CYP2S1, CYP1A2). In particular, altered expression of enzymes in the cytochrome P450 mixed-function oxidase system (CYPs) can contribute significantly to neurotoxicity by metabolizing the pesticide CPF into chlorpyrifos oxon (CPO), an active metabolite that inhibits acetylcholinesterase (Khokhar and Tyndale, 2012; Zhou et al., 2013). An increase in the levels of these scaffold proteins provides further evidence that CPF reaches the brain and likely triggers a coping response.

Despite ChE inhibition being the most plausible and common mechanism linked to behavioral impairment, there are several other mechanisms by which CPF neurotoxicity could affect cognitive function that are often overlooked [23]. Looking at the whole brain proteome of CPF-exposed and sham-treated bats, we identified enriched cellular pathways that could underlie the non-cholinergic effects of CPF on the central nervous system. For example, we detected changes in the abundance of proteins previously associated with alterations in spatial navigation (SGK1, Ma *et al.*, 2006) and associative memory (CREB, Aggleton *et al.*, 2012; PTPN5,Zhang *et al.*, 2010). Below we discuss the roles of these and other differentially abundant proteins involved in mechanisms of neurodegeneration and neuroinflammation that could explain the altered behavioral outcomes we observed [23].

Neuroinflammation contributes to the neurotoxic effects of OPs, especially after a repeated exposure at low concentrations (reviewed in Guignet and Lein, 2019). In bats exposed to CPF, we found changes in the abundance of proteins that trigger neuroinflammation through different cellular processes: I. oxidative stress (PLA2G4A, RNF112 and NOX3); II. neuronal death (PLA2G4A, SLC1A7 and SLX5), and III. regulation of pro-inflammatory mediators (PTPRT, CARMIL1, FPR2, AGTRAP and FOXP3) (See Table 1 for details). These mechanisms can also lead to neurodegeneration and synaptic disfunction in hippocampal regions important for spatial navigation and associative learning, like the CA1 (Cornu Ammonis 1) area and medial entorhinal cortex (MEC)[18,82]. Neural apoptosis in these hippocampal regions has been previously associated with impairment of spatial memory in bats exposed to imidacloprid, a neonicotinoid insecticide with a similar neurotoxic mode of action as CPF [5]. Additionally, we found that CPF exposure induced changes in proteins implicated in synaptic maintenance (PTPRT, SYX5) and neuroplasticity (PTPN5). Altogether, these mechanisms can explain the exposed bats’ deficient performance in associative memory tasks tested in the Y-maze arena.

One protein that could be particularly relevant for bat ecology is NADPH Oxidase 3 (NOX3), an enzyme highly expressed in the mammalian inner ear and the vestibular system. This protein has been positively selected in echolocating bats, suggesting its importance in spatial navigation using primarily auditory cues (reflected echoes) for perception [83]. Bats exposed to CPF showed upregulation of NOX3 after three days of exposure, which also corresponded to their worst performance in the associative memory test. Overexpression of NOX3, exacerbated by CPF exposure [84], can lead to hearing loss [85,86] and balance disorders due to the induction of reactive oxygen species (ROS), which causes neuronal loss in the central auditory system.

We have discussed some of the main protein changes that we observed and that are likely related to mechanisms of CPF neurotoxicity and impairment of behavior in bats. It is important to highlight that for some of the proteins the direction of the effects was not consistent across the two exposure durations. For some proteins, the direction of change after seven days of CPF exposure was opposite to the effect observed after three days of exposure (Fig. 8). In a multi-dose study, second and subsequent doses were found to change the biomarker responses because the system might not return to equilibrium between doses [87]. Thus, responses observed at later stages can reflect mechanisms of recovery or cellular repair rather than neurotoxic effects. On the other hand, homeostatic feedback mechanisms can prevent a sustained, unidirectional response and would tend to return proteins to baseline levels, especially when persistently altered expression can have severe consequences for cell integrity and survival. Given that we observed different protein patterns at three and seven days of exposure, and that the effects in some molecular pathways appear and disappear relatively quickly, we suggest that protein expression needs to be examined more frequently and with higher resolution to eliminate the possibility that no effect was observed when in fact changes may have occurred as an early response to exposure. In particular, late measurement when severe cellular damage has occurred may provide little mechanistic information on the toxicant mode of action [88].

While a standardized Y-maze assessment arena is an informative short-term test of learning and spatial memory, a longer-term and perhaps more ecologically relevant test for spatial navigation in bats is the ability to adopt stereotyped flight patterns when exploring novel space [31]. Studying stereotyped flight behavior provides insight into the interaction between echolocation, spatial memory, and flight control, and could help to elucidate how pesticides affect complex behavior. Stereotyped flight was found to be affected by the neonicotinoid imidacloprid, [5]. Contrary to our expectations based on previous findings, we found no difference in the consistency of or improvement in the flight trajectory over time in bats exposed to CPF (Fig. 1). Given the robust evidence of neurotoxicity that we found at the molecular level (Fig. 7, 8, and 9), and its correlation with other behavioral impairment (Fig. 2), we cannot conclude that CPF does not affect bat ability to sustain stereotyped flight. One important difference in the Hsiao et al. (2016) study is that they used newly caught bats; the relatively lower stamina of our captivity-acclimatized experimental animals could have introduced noise in the evaluation of their flight performance. Captive bats may be less motivated to initiate flight, and, in some cases, bats were unable to sustain continuous flight likely due to their increased weight in captivity, which might have influenced the accuracy of our flight trajectory estimates.

The righting reflex response is another important behavioral endpoint widely used in toxicology to assess overt neurotoxic effects of OPs pesticides. For example, rodents exposed to CPF decrease their righting performance or lose the reflex altogether [89–92]. Previous studies in bats have also found the righting reflex response to be a sensitive indicator of OPs neurotoxicity, with righting latency being positively correlated with pesticide dose [12,76,93]. Consistent with these findings, we observed a significant delay in the righting reflex response of *E. fuscus* exposed to CPF, with treated bats on average taking 50% longer to execute the maneuver compared to control group bats (Fig 5). Impairment of this reflex can have detrimental implications for survival because righting maneuvers are essential for landing—requiring bats to reorient their body and half-summersault upside down — and correcting flight trajectory after catching prey [94]. Molecular mechanisms involved in synaptoxicity could contribute to deficient cell signaling to elicit or properly coordinate this reflex response. It is important to note that while our results are consistent with previous studies in bats, the magnitude of the effect was smaller and, unlike previous studies, we did not observe other motor impairments such as the inability to fly or the presence of tremors and seizures [12,76,93]. The absence of these responses is most likely explained by the lower concentration we used to expose bats compared to previous studies that used 2-60 times greater doses.

## Conclusions

Pesticide use is a potential threat for bat populations that has been overlooked and can represent a major conservation problem in agriculture-dominant regions. Although the main mode of action of common pesticides like OPs has been widely studied, other physiological mechanisms for off-target neurotoxicity, and their consequences for individual and population health in wildlife, are still poorly understood. We found that CPF is a potent neurotoxicant that leads to behavioral impairments in bats, at low concentrations of exposure, likely due to neuronal dysfunction caused mainly by oxidative stress and neurochemical disruption. Integrating molecular and whole-animal individual responses enabled us to link neurotoxic effects with neurobehavioral outcomes. The behavioural effects we observed may compromise bat survival and lead to population declines with concomitant losses of ecological functionality and ecosystem services [3,76,95]. Our findings highlight the importance of considering endpoints other than mortality, or single molecular biomarkers when assessing the potential impact of pesticides on wildlife.

## Acknowledgments

We are very thankful to Jerrica Jamison and Raj Panchal for their assistance with the lab experiments. We thank Dr. Kathleen Delaney, Dawn Graham, and the staff of the Psychology Animal Facility at McMaster University for veterinary and animal care support.

## References

1. Boyles JG, Cryan PM, McCracken GF, Kunz TH. Economic Importance of Bats in Agriculture. Science (80-). 2011;332: 41–42. doi:10.1126/science.1201366

2. Voigt CC, Kingston T. Bats in the anthropocene. Bats Anthr Conserv Bats a Chang World. 2015; 1–9. doi:10.1007/978-3-319-25220-9_1/FIGURES/3

3. Stahlschmidt P, Bruhl CA. Bats at risk? Bat activity and insecticide residue analysis of food items in an apple orchard. Environ Toxicol Chem. 2012;31: 1556–1563. doi:10.1002/etc.1834

4. Jones G, Jacobs DS, Kunz TH, Wilig MR, Racey PA. Carpe noctem: The importance of bats as bioindicators. Endanger Species Res. 2009;8: 93–115. doi:10.3354/esr00182

5. Hsiao CJ, Lin CL, Lin TY, Wang SE, Wu CH. Imidacloprid toxicity impairs spatial memory of echolocation bats through neural apoptosis in hippocampal CA1 and medial entorhinal cortex areas. Neuroreport. 2016;27: 462–468. doi:10.1097/WNR.0000000000000562

6. Walker CH. Neurotoxic pesticides and behavioural effects upon birds. Ecotoxicology. 2003;12: 307–316. doi:10.1023/A:1022523331343

7. Grue CE, Gibert PL, Seeley ME. Neurophysiological and behavioral changes in non-target wildlife exposed to organophosphate and carbamate pesticides: Thermoregulation, food consumption, and reproduction. Am Zool. 1997;37: 369–388. doi:10.2307/3884019

8. Sandoval-Herrera N, Mena F, Espinoza M, Romero A. Neurotoxicity of organophosphate pesticides could reduce the ability of fish to escape predation under low doses of exposure. Sci Rep. 2019;9: 1–30. doi:10.1038/s41598-019-46804-6

9. Bernanke J, Köhler HR. The impact of environmental chemicals on wildlife vertebrates. Reviews of Environmental Contamination and Toxicology. New York, NY: Springer New York; 2009. pp. 1–47. doi:10.1007/978-0-387-09647-6_1

10. Naughton SX, Terry A V. Neurotoxicity in acute and repeated organophosphate exposure. Toxicology. 2018;408: 101–112. doi:10.1016/J.TOX.2018.08.011

11. Jokanović M. Neurotoxic effects of organophosphorus pesticides and possible association with neurodegenerative diseases in man: A review. Toxicology. 2018;410: 125–131. doi:10.1016/J.TOX.2018.09.009

12. Eidels RR, Sparks DW, Whitaker JO, Sprague CA. Sub-lethal Effects of Chlorpyrifos on Big Brown Bats (Eptesicus fuscus). Arch Environ Contam Toxicol. 2016;71: 322–335. doi:10.1007/s00244-016-0307-3

13. Gupta R. Toxicology of Organophosphate and Carbamate Compound. Elsevier Academic Press; 2006. Available: https://books.google.co.cr/books?hl=es&lr=&id=mFFoWG-x4rAC&oi=fnd&pg=PP1&dq=use+of+organophosphates+in+tropical+aquatic+systems&ots=g0pOuRGTTI&sig=dtDmJ7WbwxQIMI9l36qM4Z9lEUc&redir_esc=y#v=onepage&q=useoforganophosphatesintropicalaquaticsystems&f=tr

14. Kamel F, Hoppin JA. Association of pesticide exposure with neurologic dysfunction and disease. Environ Health Perspect. 2004;112: 950–958. doi:10.1289/EHP.7135

15. Moser VC. Animal models of chronic pesticide neurotoxicity. Hum Exp Toxicol. 2007;26: 321–331. doi:10.1177/0960327106072395

16. Timofeeva OA, Roegge CS, Seidler FJ, Slotkin TA, Levin ED. Persistent cognitive alterations in rats after early postnatal exposure to low doses of the organophosphate pesticide, diazinon. Neurotoxicol Teratol. 2008;30: 38–45. doi:10.1016/J.NTT.2007.10.002

17. Kohler H-R, Triebskorn R. Wildlife Ecotoxicology of Pesticides: Can We Track Effects to the Population Level and Beyond? Science (80-). 2013;341: 759–765. doi:10.1126/science.1237591

18. Yovel Y, Ulanovsky N. Bat navigation. In: Byrne JH, editor. Learning and Memory: A Comprehensive Reference, Second Edition. Elsevier; 2017. pp. 1–2402.

19. Keifer MC, Firestone J. Neurotoxicity of Pesticides. J Agromedicine. 2007;12: 17–25. doi:10.1300/J096v12n01_03

20. Amiard-Triquet C, Amiard JC. Ecological biomarkers: indicators of ecotoxicological effects. CRC press. FL; 2012. doi:10.1017/CBO9781107415324.004

21. Lionetto MG, Caricato R, Calisi A, Giordano ME, Schettino T. Acetylcholinesterase as a biomarker in environmental and occupational medicine: New insights and future perspectives. Biomed Res Int. 2013;2013. doi:10.1155/2013/321213

22. Jebali J, Khedher S Ben, Sabbagh M, Kamel N, Banni M, Boussetta H. Cholinesterase activity as biomarker of neurotoxicity: utility in the assessment of aquatic environment contamination. Rev Gestão Costeira Integr. 2013;13: 525–537. doi:10.5894/rgci430

23. Tsai YH, Lein PJ. Mechanisms of organophosphate neurotoxicity. Curr Opin Toxicol. 2021;26: 49–60. doi:10.1016/J.COTOX.2021.04.002

24. Garcia-Reyero N, Perkins EJ. Systems biology: Leading the revolution in ecotoxicology. Environmental Toxicology and Chemistry. 2011. pp. 265–273. doi:10.1002/etc.401

25. Saper CB, Scammell TE, Lu J. Hypothalamic regulation of sleep and circadian rhythms. Nature. 2005;437: 1257–1263. doi:10.1038/nature04284

26. Liang X, Martyniuk CJ, Simmons DBD. Are we forgetting the “proteomics” in multi-omics ecotoxicology? Comp Biochem Physiol Part D Genomics Proteomics. 2020;36: 100751. doi:10.1016/J.CBD.2020.100751

27. Valcu CM, Kempenaers B. Proteomics in behavioral ecology. Behav Ecol. 2015;26: 1–15. doi:10.1093/BEHECO/ARU096

28. US EPA. Guidelines for Exposure Assessment. Risk Assess Forum. 1992;57: 22888–22938. doi:EPA/600/Z-92/001

29. Solomon KR, Giesy JP, Kendall RJ, Best LB, Coats JR, Dixon KR, et al. Chlorpyrifos: Ecotoxicological risk assessment for birds and mammals in corn agroecosystems. Hum Ecol Risk Assess. 2001;7: 497–632. doi:10.1080/20018091094510

30. Barchi JR, Knowles JM, Simmons JA. Spatial memory and stereotypy of flight paths by big brown bats in cluttered surroundings. J Exp Biol. 2013;216: 1053–1063. doi:10.1242/jeb.073197

31. Barchi JR, Knowles JM, Simmons JA. Spatial memory and stereotypy of flight paths by big brown bats in cluttered surroundings. J Exp Biol. 2013;216: 1053–1063. doi:10.1242/jeb.073197

32. Heredia-López FJ, Álvarez-Cervera FJ, Collí-Alfaro JG, Bata-García JL, Arankowsky-Sandoval G, Góngora-Alfaro JL. An automated Y-maze based on a reduced instruction set computer (RISC) microcontroller for the assessment of continuous spontaneous alternation in rats. Behav Res Methods. 2016;48: 1631–1643. doi:10.3758/S13428-015-0674-0/TABLES/2

33. Cleal M, Fontana BD, Ranson DC, McBride SD, Swinny JD, Redhead ES, et al. The Free-movement pattern Y-maze: A cross-species measure of working memory and executive function. Behav Res Methods 2020 532. 2020;53: 536–557. doi:10.3758/S13428-020-01452-X

34. Marcos Gorresen P, Cryan PM, Dalton DC, Wolf S, Bonaccorso FJ. Ultraviolet vision may be widespread in bats. Acta Chiropterologica. 2015;17: 193–198. doi:10.3161/15081109ACC2015.17.1.017

35. De Fanis E, Jones G. The role of odour in the discrimination of conspecifics by pipistrelle bats. A&l Behav. 1995;49: 835–839.

36. Bartonička TÁ, Kaňuch P, Bímova B, Bryja J. Olfactory discrimination between two cryptic species of bats Pipistrellus pipistrellus and P. pygmaeus. https://doi.org/1025225/fozo.v59.i3.a22010. 2010;59: 175–182. doi:10.25225/FOZO.V59.I3.A2.2010

37. Kilgour RJ, Faure PA, Brigham RM. Evidence of social preferences in big brown bats (Eptesicus fuscus). https://doi.org/101139/cjz-2013-0057. 2013;91: 756–760. doi:10.1139/CJZ-2013-0057

38. Greville LJS, Tam AG, Faure PA. Evaluating odour and urinary sex preferences in the big brown bat (Eptesicus fuscus). Can J Zool. 2021;99: 930–938. doi:10.1139/cjz-2021-0067

39. Riskin DK, Parsons S, Schutt WA, Carter GG, Hermanson JW. Terrestrial locomotion of the New Zealand short-tailed bat Mystacina tuberculata and the common vampire bat Desmodus rotundus. J Exp Biol. 2006;209: 1725–1736. doi:10.1242/JEB.02186

40. Ellman G, Courtney K, Andres V. A new and rapid colorimetric determination of acetylcholinesterase activity. Biochemical. 1961 [cited 22 Oct 2016]. Available: http://www.sciencedirect.com/science/article/pii/0006295261901459

41. Bradford MM. A rapid and sensitive method for the quantitation of microgram quantities of protein utilizing the principle of protein-dye binding. Anal Biochem. 1976;72: 248–54. Available: http://www.ncbi.nlm.nih.gov/pubmed/942051

42. Bates D, Mächler M, Bolker BM, Walker SC. Fitting Linear Mixed-Effects Models Using lme4. J Stat Softw. 2015;67: 1–48. doi:10.18637/JSS.V067.I01

43. R Core Team. R: A language and environment for statistical computing. Viena, Austria: R Foundation for Statistical Computing; 2019.

44. Chong J, Wishart DS, Xia J. Using MetaboAnalyst 4.0 for Comprehensive and Integrative Metabolomics Data Analysis. Curr Protoc Bioinforma. 2019;68: e86. doi:10.1002/CPBI.86

45. Supek F, Bošnjak M, Škunca N, Šmuc T. REVIGO Summarizes and Visualizes Long Lists of Gene Ontology Terms. PLoS One. 2011;6: e21800. doi:10.1371/JOURNAL.PONE.0021800

46. Sharma P, Nag A. CUL4A ubiquitin ligase: a promising drug target for cancer and other human diseases. Open Biol. 2014;4. doi:10.1098/RSOB.130217

47. Li T, Hung MS, Wang Y, Mao JH, Tan JL, Jahan K, et al. Transgenic mice for cre-inducible overexpression of the Cul4A gene. Genesis. 2011;49: 134–141. doi:10.1002/DVG.20708

48. Stamou M, Wu X, Kania-Korwel I, Lehmler HJ, Lein PJ. Cytochrome P450 mRNA expression in the rodent brain: Species-, sex-, and region-dependent differences. Drug Metab Dispos. 2014;42: 239–244. doi:10.1124/DMD.113.054239/-/DC1

49. McMillan DM, Tyndale RF. CYP-mediated drug metabolism in the brain impacts drug response. Pharmacol Ther. 2018;184: 189–200. doi:10.1016/J.PHARMTHERA.2017.10.008

50. Patel SL, Prakash J, Gupta V. Decreased mRNA expression level of FOXP3 correlate with TNF-α in peripheral blood mononuclear cells (PBMCs) from rheumatoid arthritis patients: A case control study. Curr Orthop Pract. 2022;33: 73–80. doi:10.1097/BCO.0000000000001067

51. Ramos JSA, Pedroso TMA, Godoy FR, Batista RE, de Almeida FB, Francelin C, et al. Multi-biomarker responses to pesticides in an agricultural population from Central Brazil. Sci Total Environ. 2021;754: 141893. doi:10.1016/J.SCITOTENV.2020.141893

52. Kalpachidou T, Makrygiannis AK, Pavlakis E, Stylianopoulou F, Chalepakis G, Stamatakis A. Behavioural effects of extracellular matrix protein Fras1 depletion in the mouse. Eur J Neurosci. 2021;53: 3905–3919. doi:10.1111/ejn.14759

53. Short K, Wiradjaja F, Smyth I. Let’s stick together: The role of the Fras1 and Frem proteins in epidermal adhesion. IUBMB Life. 2007;59: 427–435. doi:10.1080/15216540701510581

54. Dasari S, Swamy GANJAYI M, Meriga B. Glutathione S-transferase is a good biomarker in acrylamide induced neurotoxicity and genotoxicity. 2018. doi:10.2478/intox-2018-0007

55. Björk K, Saarikoski ST, Arlinde C, Kovanen L, Osei‐Hyiaman D, Ubaldi M, et al. Glutathione-S-transferase expression in the brain: possible role in ethanol preference and longevity. FASEB J. 2006;20: 1826–1835. doi:10.1096/FJ.06-5896COM

56. Cooney SJ, Bermudez-Sabogal SL, Byrnes KR. Cellular and temporal expression of NADPH oxidase (NOX) isotypes after brain injury. J Neuroinflammation. 2013;10: 155. doi:10.1186/1742-2094-10-155

57. Nakano Y, Banfi B, Jesaitis AJ, Dinauer MC, Allen LAH, Nauseef WM. Critical roles for p22phox in the structural maturation and subcellular targeting of Nox3. Biochem J. 2007;403: 97–108. doi:10.1042/BJ20060819

58. Lee J-R. Protein tyrosine phosphatase PTPRT as a regulator of synaptic formation and neuronal development. BMB Rep. 2015;48: 249. doi:10.5483/BMBREP.2015.48.5.037

59. Dattilo V, Amato R, Perrotti N, Gennarelli M. The Emerging Role of SGK1 (Serum- and Glucocorticoid-Regulated Kinase 1) in Major Depressive Disorder: Hypothesis and Mechanisms. Front Genet. 2020;11: 826. doi:10.3389/FGENE.2020.00826/BIBTEX

60. Margiotta A. Role of SNAREs in Neurodegenerative Diseases. Cells. 2021;10. doi:10.3390/CELLS10050991

61. PT L, CV H, MT B, G van den B. Stx5-Mediated ER-Golgi Transport in Mammals and Yeast. Cells. 2019;8: 780. doi:10.3390/CELLS8080780

62. Lee J-A, Gao F-B. Neuronal Functions of ESCRTs. Exp Neurobiol. 2012;21: 9. doi:10.5607/EN.2012.21.1.9

63. Loera-Valencia R, Eroli F, Garcia-Ptacek S, Maioli S. Brain Renin–Angiotensin System as Novel and Potential Therapeutic Target for Alzheimer’s Disease. Int J Mol Sci 2021, Vol 22, Page 10139. 2021;22: 10139. doi:10.3390/IJMS221810139

64. Stark BC, Cooper JA. Differential expression of CARMIL-family genes during zebrafish development. Cytoskeleton (Hoboken). 2015;72: 534. doi:10.1002/CM.21257

65. Cha J, Jeon T-W, Lee CG, Oh ST, Yang H-B, Choi K-J, et al. International Journal of Hyperthermia Electro-hyperthermia inhibits glioma tumorigenicity through the induction of E2F1-mediated apoptosis Electro-hyperthermia inhibits glioma tumorigenicity through the induction of E2F1-mediated apoptosis. Int J Hyperth. 2015;31: 784. doi:10.3109/02656736.2015.1069411

66. Deng Y, Zhang Y, Lu Y, Zhao Y, Ren H. Hepatotoxicity and nephrotoxicity induced by the chlorpyrifos and chlorpyrifos-methyl metabolite, 3,5,6-trichloro-2-pyridinol, in orally exposed mice. Sci Total Environ. 2016;544: 507–514. doi:10.1016/J.SCITOTENV.2015.11.162

67. Mao W, Guo Z, Dai Y, Nie J, Li B, Pei H, et al. LNC CRYBG3 inhibits tumor growth by inducing M phase arrest. J Cancer. 2019;10: 2764–2770. doi:10.7150/JCA.31703

68. Krovi SH, Zhang J, Michaels-Foster MJ, Brunetti T, Loh L, Scott-Browne J, et al. Single-cell RNA transcriptomics identifies Hivep3 as essential in regulating the development of innate-like T lymphocytes. bioRxiv. 2020; 2020.06.08.135129. doi:10.1101/2020.06.08.135129

69. Hicar MD, Liu Y, Allen CE, Wu LC. Structure of the human zinc finger protein HIVEP3: molecular cloning, expression, exon-intron structure, and comparison with paralogous genes HIVEP1 and HIVEP2. Genomics. 2001;71: 89–100. doi:10.1006/GENO.2000.6425

70. Sarkar C, Jones JW, Hegdekar N, Thayer JA, Kumar A, Faden AI, et al. PLA2G4A/cPLA2-mediated lysosomal membrane damage leads to inhibition of autophagy and neurodegeneration after brain trauma. Autophagy. 2020;16: 466. doi:10.1080/15548627.2019.1628538

71. Law MH, Cotton RGH, Berger GE. The role of phospholipases A2 in schizophrenia. Mol Psychiatry 2006 116. 2006;11: 547–556. doi:10.1038/sj.mp.4001819

72. Yang C-H, Huang C-C, Hsu K-S. A Critical Role for Protein Tyrosine Phosphatase Nonreceptor Type 5 in Determining Individual Susceptibility to Develop Stress-Related Cognitive and Morphological Changes. J Neurosci. 2012;32: 7550. doi:10.1523/JNEUROSCI.5902-11.2012

73. Fitzpatrick CJ, Lombroso PJ. The role of striatal-enriched protein tyrosine phosphatase (STEP) in cognition. Front Neuroanat. 2011;0: 47. doi:10.3389/FNANA.2011.00047/BIBTEX

74. Tsou JH, Yang YC, Pao PC, Lin HC, Huang NK, Lin ST, et al. Important Roles of Ring Finger Protein 112 in Embryonic Vascular Development and Brain Functions. Mol Neurobiol. 2017;54: 2286–2300. doi:10.1007/S12035-016-9812-7

75. Zhou Y, Danbolt NC. GABA and glutamate transporters in brain. Front Endocrinol (Lausanne). 2013;4: 165. doi:10.3389/FENDO.2013.00165/BIBTEX

76. Clark DR. Toxicity of methyl parathion to bats: Mortality and coordination loss. Environ Toxicol Chem. 1986;5: 191–195. doi:10.1002/etc.5620050210

77. Trudeau S, Cartier GS. Biochemical Methods to Determine Cholinesterase Activity in Wildlife Exposed to Pesticides. Technical Report Series Number 338. 2000.

78. Ma YL, Tsai MC, Hsu WL, Lee EHY. SGK protein kinase facilitates the expression of long-term potentiation in hippocampal neurons. Learn Mem. 2006;13: 114–118. doi:10.1101/lm.179206

79. Aggleton JP, Brown MW, Albasser MM. Contrasting brain activity patterns for item recognition memory and associative recognition memory: Insights from immediate-early gene functional imaging. Neuropsychologia. 2012;50: 3141–3155. doi:10.1016/J.NEUROPSYCHOLOGIA.2012.05.018

80. Zhang Y, Kurup P, Xu J, Carty N, Fernandez SM, Nygaard HB, et al. Genetic reduction of striatal-enriched tyrosine phosphatase (STEP) reverses cognitive andcellular deficits in an Alzheimer’s disease mouse model. Proc Natl Acad Sci U S A. 2010;107: 19014–19019. doi:10.1073/PNAS.1013543107/-/DCSUPPLEMENTAL

81. Guignet M, Lein PJ. Neuroinflammation in organophosphate-induced neurotoxicity. Adv Neurotoxicology. 2019;3: 35–79. doi:10.1016/BS.ANT.2018.10.003

82. Wilson MA, McNaughton BL. Dynamics of the Hippocampal Ensemble Code for Space. Science (80-). 1993;261: 1055–1058. doi:10.1126/SCIENCE.8351520

83. Seim I, Fang X, Xiong Z, Lobanov A V., Huang Z, Ma S, et al. Genome analysis reveals insights into physiology and longevity of the Brandt’s bat Myotis brandtii. Nat Commun 2013 41. 2013;4: 1–8. doi:10.1038/ncomms3212

84. Dominah GA, McMinimy RA, Kallon S, Kwakye GF. Acute exposure to chlorpyrifos caused NADPH oxidase mediated oxidative stress and neurotoxicity in a striatal cell model of Huntington’s disease. Neurotoxicology. 2017;60: 54–69. doi:10.1016/J.NEURO.2017.03.004

85. Bánfi B, Malgrange B, Knisz J, Steger K, Dubois-Dauphin M, Krause KH. NOX3, a Superoxide-generating NADPH Oxidase of the Inner Ear. J Biol Chem. 2004;279: 46065–46072. doi:10.1074/JBC.M403046200

86. Mohri H, Ninoyu Y, Sakaguchi H, Hirano S, Saito N, Ueyama T. Nox3-Derived Superoxide in Cochleae Induces Sensorineural Hearing Loss. J Neurosci. 2021;41: 4716–4731. doi:10.1523/JNEUROSCI.2672-20.2021

87. Nicholson JK, Connelly J, Lindon JC, Holmes E. Metabonomics: a platform for studying drug toxicity and gene function. Nat Rev Drug Discov 2002 12. 2002;1: 153–161. doi:10.1038/nrd728

88. Dam K, Seidler FJ, Slotkin TA. Transcriptional biomarkers distinguish between vulnerable periods for developmental neurotoxicity of chlorpyrifos: Implications for toxicogenomics. Brain Res Bull. 2003;59: 261–265. doi:10.1016/S0361-9230(02)00874-2

89. Chanda SM, Harp P, Liu J, Pope CN. Comparative developmental and maternal neurotoxicity following acute gestational exposure to chlorpyrifos in rats. J Toxicol Environ Health. 1995;44: 189–202. doi:10.1080/15287399509531954

90. Chanda SM, Pope CN. Neurochemical and neurobehavioral effects of repeated gestational exposure to chlorpyrifos in maternal and developing rats. Pharmacol Biochem Behav. 1996;53: 771–776. doi:10.1016/0091-3057(95)02105-1

91. Ambali SF, Ayo JO. Vitamin C Attenuates Chronic Chlorpyrifos-induced Alteration of Neurobehavioral Parameters in Wistar Rats. Toxicol Int. 2012;19: 144. doi:10.4103/0971-6580.97211

92. Khokhar JY, Tyndale RF. Rat Brain CYP2B-Enzymatic Activation of Chlorpyrifos to the Oxon Mediates Cholinergic Neurotoxicity. Toxicol Sci. 2012;126: 325–335. doi:10.1093/TOXSCI/KFS029

93. Clark D, Rattner B. Orthene toxicity to little brown bats (Myotis lucifugus): acetylcholinesterase inhibition, coordination loss, and mortality. Environ Toxicol Chem. 1987;6: 705–708. doi:10.1016/0032-5910(72)80020-2

94. Bergou AJ, Swartz SM, Vejdani H, Riskin DK, Reimnitz L, Taubin G, et al. Falling with Style: Bats Perform Complex Aerial Rotations by Adjusting Wing Inertia. PLOS Biol. 2015;13: e1002297. doi:10.1371/JOURNAL.PBIO.1002297

95. Jefferies DJ. Organochlorine insecticide residues in British bats and their significance. J Zool. 2009;166: 245–263. doi:10.1111/j.1469-7998.1972.tb04088.x

