## Supplementary table 1 for "Common insecticide affects spatial navigation in bats at environmentally-realistic doses"

Supplementary table 1: Description of differentially abundant proteins in brain tissue of *Eptesicus fuscus* exposed to CPF.

| Uniprot ID | Protein ID | Protein name | Protein class/molecular function | ANOVA P-value | B-H P-value |
|---|---|---|---|---|---|
| **Q13619** | CUL4A | Cullin-4A | Ubiquitin-protein ligase | 0.011753 | 0.692544 |
| **Q5T3I0** | GPATCH4 | G patch domain-containing protein 4 | RNA metabolism protein | 0.012687 | 0.692544 |
| **Q5VZK9** | CARMIL1 | F-actin-uncapping protein LRRC16A | Protein-binding activity modulator | 0.014469 | 0.692544 |
| **Q9HBY0** | NOX3 | NADPH oxidase 3 | Oxidase | 0.015165 | 0.692544 |
| **O14522** | PTPRT | Receptor-type tyrosine-protein phosphatase T | Protein phosphatase | 0.01863 | 0.692544 |
| **Q9H2J7** | SLC6A15 | Sodium-dependent neutral amino acid transporter B(0)AT2 | Primary active transporter | 0.022254 | 0.710485 |
| **P54829** | PTPN5 | Tyrosine-protein phosphatase non-receptor type 5 | Phosphotyrosine residue binding | 0.022442 | 0.692544 |
| **P10253** | GAA | Lysosomal alpha-glucosidase | Glucosidase | 0.023466 | 0.710485 |
| **Q4G0M1** | ERFE | Erythroferrone | Peptide hormone | 0.026973 | 0.692544 |
| **Q68DQ2** | CRYBG3 | Very large A-kinase anchor protein | Structural protein | 0.027556 | 0.692544 |
| **Q99676** | ZNF184 | Zinc finger protein 184 | C2H2 zinc finger transcription factor | 0.029139 | 0.692544 |
| **Q460N5** | PARP14 | Protein mono-ADP-ribosyltransferase PARP14 | ADP-ribosyltransferase | 0.029989 | 0.692544 |
| **Q5TZJ5** | SPATA31A1 | Spermatogenesis-associated protein 31A1 | Cell differentiation | 0.030058 | 0.692544 |
| **O95727** | CRTAM | Cytotoxic and regulatory T-cell molecule | Cell differentiation | 0.031015 | 0.692544 |
| **O15379** | HDAC3 | Histone deacetylase 3 | Histone modifying enzyme | 0.031894 | 0.692544 |
| **Q5T1R4** | HIVEP3 | Transcription factor HIVEP3 | Transcription factor | 0.033405 | 0.710485 |
| **Q7Z591** | AKNA | Microtubule organization protein AKNA | DNA binding | 0.038401 | 0.710485 |
| **P47712** | PLA2G4A | Cytosolic phospholipase A2 | Phospholipase | 0.04137 | 0.710485 |
| **Q9NX78** | TMEM260 | Transmembrane protein 260 | Transmembrane protein | 0.041748 | 0.692544 |
| **Q8NHU2** | CFAP61 | Cilia- and flagella-associated protein 61 | Structural protein | 0.043203 | 0.692544 |
| **Q86SQ0** | PHLDB2 | Pleckstrin homology-like domain family B member 2 | Cadherin binding | 0.04544 | 0.710485 |
| **Q92859** | NEO1 | Neogenin | Cell adhesion molecule | 0.048292 | 0.692544 |
| **P04080** | CYTB | Cystatin-B | Protease inhibitor | 0.049246 | 0.692544 |
| **O75962** | TRIO | Triple functional domain protein | Guanyl-nucleotide exchange factor | 0.0495 | 0.710485 |
| **O75366** | AVIL | Advillin | Non-motor actin binding protein | 0.049674 | 0.692544 |
| **Q96SQ9** | CYP2S1 | Cytochrome P450 2S1 | Oxygenase | 0.050022 | 0.710485 |
| **Q9P2I0** | CPSF2 | Cleavage and polyadenylation specificity factor subunit 2 | RNA binding | 0.050249 | 0.692544 |
| **Q6RW13** | AGTRAP | Type-1 angiotensin II receptor-associated protein | Receptor-mediated signaling | 0.05039 | 0.692544 |
| **Q9Y6Y1** | CAMTA1 | Calmodulin-binding transcription activator 1 | DNA-binding transcription factor | 0.051754 | 0.692544 |
| **Q99797** | MIPEP | Mitochondrial intermediate peptidase | Metalloprotease | 0.052685 | 0.692544 |
| **Q86UN2** | RTN4RL1 | Reticulon-4 receptor-like 1 | Transmembrane signal receptor | 0.056031 | 0.692544 |
| **O00341** | SLC1A7 | Excitatory amino acid transporter 5 | Primary active transporter | 0.056759 | 0.710485 |

| | | | | | |
|---|---|---|---|---|---|
| **O60522** | TDRD6 | Tudor domain-containing protein 6 | Scaffold/adaptor protein | 0.057422 | 0.710485 |
| **Q86XT2** | VPS37D | Vacuolar protein sorting-associated protein 37D | Membrane trafficking regulatory protein | 0.060499 | 0.692544 |
| **Q13061** | TRDN | Triadin | Signaling receptor binding | 0.066839 | 0.692544 |
| **Q86XX4** | FRAS1 | Extracellular matrix organizing protein FRAS1 | Extracellular matrix protein | 0.068529 | 0.692544 |
| **Q14146** | URB2 | Unhealthy ribosome biogenesis protein 2 homolog | Regulation of signal transduction | 0.071511 | 0.692544 |
| **P0CG30** | GSTT2 | Glutathione S-transferase theta-2B | Transferase | 0.072609 | 0.710485 |
| **Q6PCB0** | VWA1 | von Willebrand factor A domain-containing protein 1 | Extracellular matrix structural protein | 0.073997 | 0.692544 |
| **O75509** | TNFRSF21 | Tumor necrosis factor receptor superfamily member 21 | Transmembrane signal receptor | 0.074017 | 0.710485 |
| **Q13190** | STX5 | Syntaxin-5 | SNARE protein | 0.076432 | 0.692544 |
| **Q5VWM4** | PRAMEF8 | PRAME family member 8 | Retinoic acid receptor binding | 0.077198 | 0.710485 |
| **Q14929** | ZNF169 | Zinc finger protein 169 | C2H2 zinc finger transcription factor | 0.077373 | 0.710485 |
| **Q96I99** | SUCLG2 | Succinate--CoA ligase [GDP-forming] subunit beta, mitochondrial | Ligase | 0.078623 | 0.692544 |
| **P25090** | FPR2 | N-formyl peptide receptor 2 | G-protein coupled receptor | 0.082468 | 0.692544 |
| **Q92925** | SMARCD2 | SWI/SNF-related matrix-associated actin-dependent regulator of chromatin subfamily D member 2 | Chromatin/chromatin-binding, or -regulatory protein | 0.084272 | 0.710485 |
| **Q96KJ9** | COX4I2 | Cytochrome c oxidase subunit 4 isoform 2, mitochondrial | Oxidase | 0.084963 | 0.710485 |
| **Q96LB1** | MRGPRX2 | Mas-related G-protein coupled receptor member X2 | G-protein coupled receptor | 0.089249 | 0.692544 |
| **P02730** | SLC4A1 | Band 3 anion transport protein | Secondary carrier transporter | 0.089453 | 0.710485 |
| **Q3KPI0** | CEACAM21 | Carcinoembryonic antigen-related cell adhesion molecule 21 | Immunoglobulin receptor superfamily | 0.093556 | 0.708194 |
| **Q9Y6R7** | FCGBP | IgGFc-binding protein | Extracellular matrix protein | 0.094027 | 0.710485 |
| **Q8TEX9** | IPO4 | Importin-4 | Transporter | 0.097561 | 0.710485 |
| **Q9UKN5** | PRDM4 | PR domain zinc finger protein 4 | C2H2 zinc finger transcription factor | 0.1017 | 0.692544 |

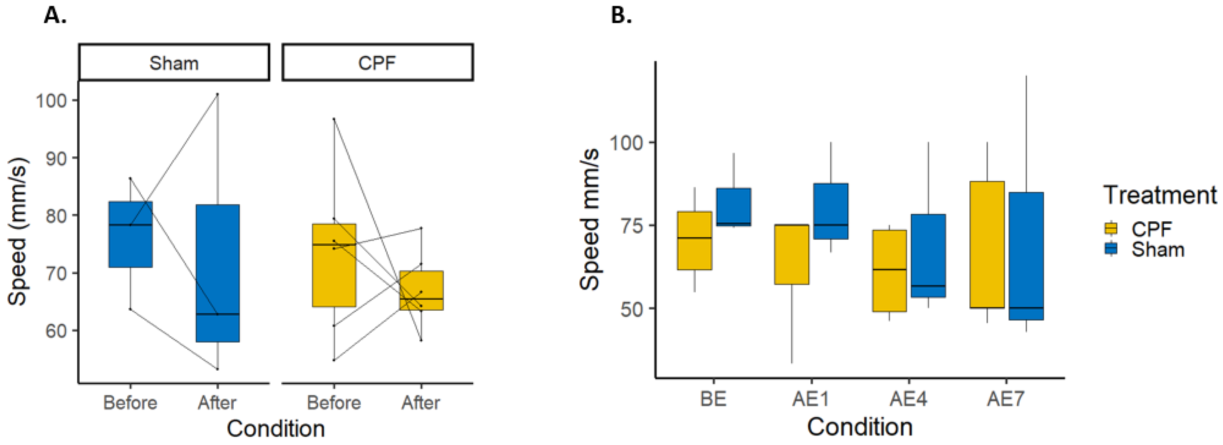


Supplementary figure 1. Cognitive behavior task performance of *E. fuscus* in Y-maze testing for sham-treated (blue) and bats exposed to CPF for seven days (yellow). A. Average crawling speed before and after treatment.Thin lines in top panel connect data for the same individual; B. Average crawling speed before exposure (BE) and 1, 3 and 7 days after exposure (i.e., AE1, AE3 and AE7).
